# The nuclear Argonaute HRDE-1 directs target gene re-localization and shuttles to nuage to promote small RNA mediated inherited silencing

**DOI:** 10.1101/2022.10.04.510877

**Authors:** Yue-He Ding, Humberto Ochoa, Takao Ishidate, Craig C. Mello

## Abstract

Argonaute small-RNA pathways engage heterochromatin-silencing co-factors to promote transgenerational inheritance in animals. However, little is known about how heterochromatin and small-RNA pathways interact to transmit silencing. Here we show that the induction of heterochromatin silencing in C. elegans by RNAi or by artificially tethering pathway components to target RNA correlates with the co-localization of the target alleles in pachytene nuclei. Tethering the nuclear Argonaute WAGO-9/HRDE-1 induces heterochromatin formation, but also functions independently to induce small-RNA amplification. We show that HRDE-1 shuttles to nuage domains called mutator foci where amplification is thought to occur. Tethering a heterochromatin-silencing factor, NRDE-2, induces heterochromatin silencing and also induces the de-novo synthesis of HRDE-1 guide RNAs, and through HRDE-1 acts to further amplify downstream small-RNA silencing. Our findings support a model in which HRDE-1 functions both upstream, to initiate heterochromatin silencing, and downstream, to stimulate small-RNA amplification, establishing a self-enforcing mechanism that propagates silencing to offspring.

## Introduction

In many animal germlines, small-RNA/Argonaute pathways function transgenerationally to install and reinforce chromatin silencing essential for fertility. For example, in flies, worms and mammals, members of the PIWI Argonaute family engage genomically encoded small RNAs termed PIWI-interacting RNAs (piRNAs) that silence transposons to maintain genome integrity (Kasschau et al., 2007; Czech et al., 2008; Ghildiyal et al., 2008; Tam et al., 2008; Watanabe et al., 2008; Gu et al., 2009). Although the details differ, all transgenerational small RNA silencing pathways studied to date require amplification and engagement of secondary Argonautes (Weick and Miska, 2014). Many of the components of the amplification machinery localize prominently in peri-nuclear non-membranous organelles called nuage. However, how the amplification system in nuage communicates with and drives the nuclear events during the initiation and maintenance of transgenerational silencing is not well understood.

In *C. elegans*, transgenerational silencing can be initiated by the PIWI pathway, by the canonical dsRNA-induced RNAi pathway, or by intronless mRNA (Fire et al., 1998; Ashe et al., 2012; Shirayama et al., 2012; Makeyeva et al., 2021). Inherited silencing is maintained by a family of related downstream worm-specific Argonautes (WAGO Argonautes) guided by small RNAs (22G-RNAs) produced by cellular RNA-dependent RNA polymerase. Once established, inherited silencing can be propagated independently of the initiating cues via continuous cycles of WAGO 22G-RNA amplification and transmission of the WAGO Argonautes and their small RNA co-factors to progeny (Lee et al., 2012; Shirayama et al., 2012; Sapetschnig et al., 2015; Spracklin et al., 2017).

The nuclear WAGO Argonaute, HRDE-1/WAGO-9, plays a central role in transgenerational silencing in *C. elegans* (Buckley et al., 2012; Rechavi et al., 2014). HRDE-1 is thought to engage nascent transcripts at target loci to induce heterochromatin and transcriptional silencing through the nuclear RNAi pathway (Buckley et al., 2012; Almeida et al., 2019). HRDE-1 promotes the transgenerational silencing of many genes (Kim et al., 2021) and is thought to do so by recruiting chromatin remodeling factors, including nucleosome remodeling and deacetylase complex (NuRD) and histone methyltransferases (e.g., MET-2, SET-25, SET-32) (Ashe et al., 2012; Towbin et al., 2012; Kim et al., 2021). The nuclear RNAi pathway is also required for the spreading of secondary small RNAs from piRNA target sites (Luteijn et al., 2012; Sapetschnig et al., 2015).

Transgenerational silencing requires a series of events that are thought to occur in the nuage, nucleus and cytoplasm. Because all of these events are essential for the cycle of inherited silencing their order has been difficult to determine. For example, it is not known whether the nuclear Argonaute HRDE-1 can trigger RdRP recruitment and amplification of small RNAs directly or whether it must instead act through the induction of heterochromatin at its targets to elicit small RNA amplification. Here we use the phage lambda N (λN)-boxB tethering system (Baron-Benhamou et al., 2004; Bühler et al., 2006; Wedeles et al., 2013; Aoki et al., 2021; Cornes et al., 2022) to recruit (tether) HRDE-1 or the nuclear silencing factor NRDE-2 to a reporter mRNA. In principle, tethering enables initiation of silencing in the absence of upstream initiators such as piRNAs or dsRNA and with appropriate genetic tests can be used to order events in the pathway. We show that tethering either HRDE-1 or NRDE-2 can induce a complete silencing response, including small-RNA amplification and transgenerational silencing that persists even after the λN-fusion protein is crossed from the strain. Tethering NRDE-2 initiates chromatin silencing through *nrde-4* and independently of *hrde-1* but requires *hrde-1* for small RNA amplification. By contrast, tethering HRDE-1 stimulates chromatin silencing through NRDE-2 and NRDE-4 but can elicit small-RNA amplification independently of both these chromatin-silencing factors. Mutants expected to abolish small RNA binding by HRDE-1 disarm silencing and cause HRDE-1 to become cytoplasmic. Tethering these mutant HRDE-1 proteins failed to initiate heterochromatin formation, but strongly initiate silencing that depends on amplification of small RNAs proximal to the tether site. The HRDE-1 amino acid sequences required to recruit the small-RNA amplification machinery reside in the N-terminal half of the protein (the N-Terminal Domain, NTD). The HRDE-1 NTD protein—like full-length HRDE-1—co-localized with MUT-16 in subdomains of cytoplasmic nuage (Mutator foci), where small-RNA amplification factors reside (Phillips et al., 2012). Our findings suggest that HRDE-1 lies at a nexus in the silencing pathway, shuttling from the nucleus to the nuage and back, to coordinate the nuclear and cytoplasmic events of transgenerational silencing.

## Results

### HRDE-1 and NRDE-2 tethering induces transgenerational silencing

To order events in inherited silencing, we sought to uncouple initiation and maintenance of silencing. To do this we used the phage λN-box-B tethering system to recruit nuclear silencing factors HRDE-1 or NRDE-2 to a target reporter that is robustly expressed in the germline (Figure 1A). We hypothesized that if artificial recruitment of a silencing factor mimics a physiological event, then it should elicit a silencing response that is independent of upstream factors but depends on known downstream factors. For example, directly tethering a chromatin factor should, in principle, induce silencing without requiring machinery necessary to amplify the small RNAs that would normally guide the chromatin silencing machinery to the appropriate targets.

**Figure 1.**
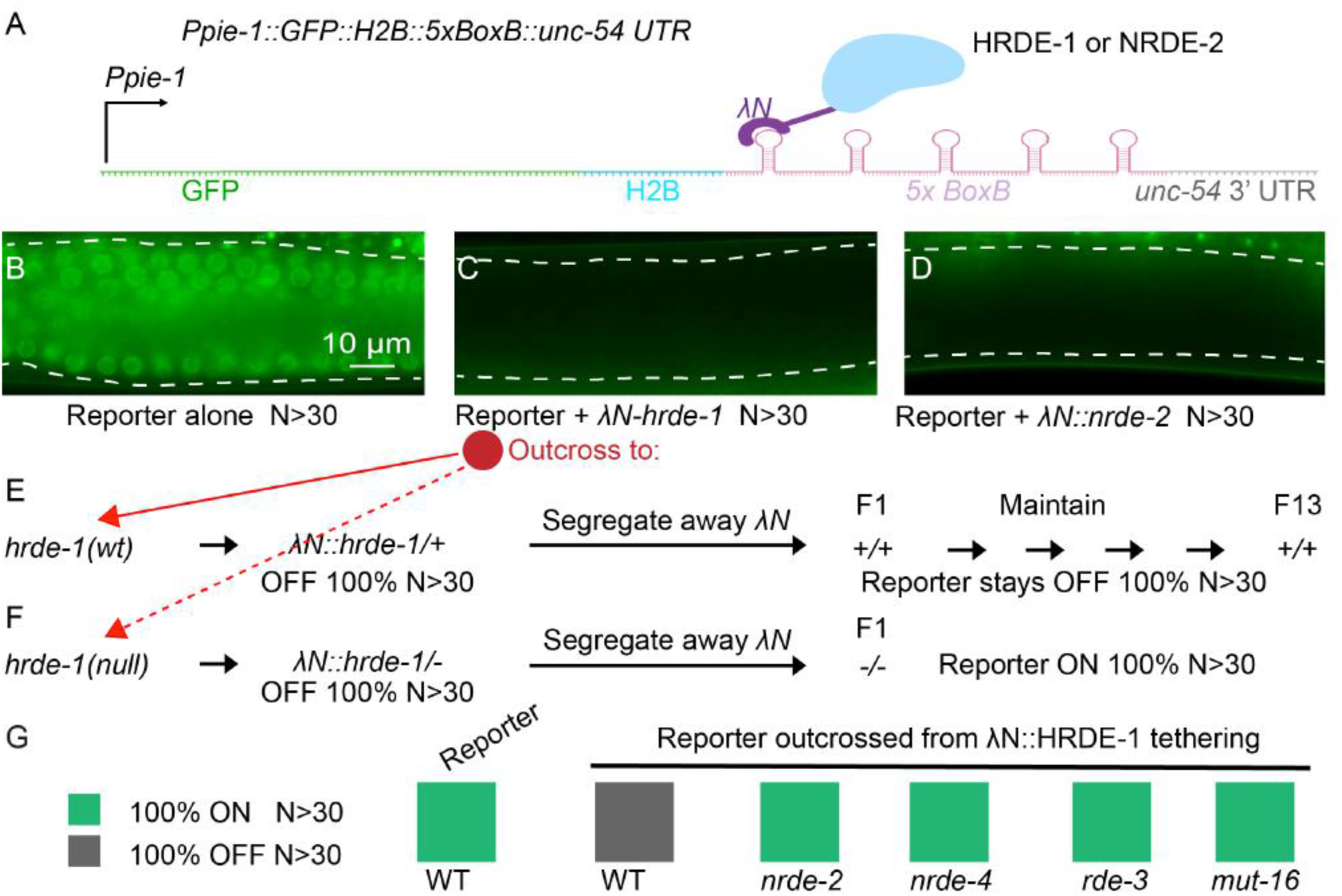
HRDE-1 tethering caused reporter silencing and generated the silencing memory. (A) Scheme of λN-BoxB tethering system. A sequence encoding five BoxB hairpins (*5xBoxB*) was inserted immediately after the coding region of the *GFP::his-58(H2B)* transgene and before the *unc-54* 3’ UTR. The reporter is driven by the *pie-1* promoter (*Ppie-1*). The BoxB sites recruit λN::HRDE-1 or λN::NRDE-2 fusion proteins thereby tethering HRDE-1 or NRDE-2 to the reporter RNA. (B) Representative fluorescence image of a syncytial germline (outlined by dashed lines) in the absence of tethering. The image represents 100% of worms scored, N>30. (C) Representative fluorescence image in the presence of HRDE-1 tethering. The image represents 100% of worms scored, N>30. (D) Representative fluorescence image in the presence of NRDE-2 tethering. The image represents 100% of worms scored, N>30. (E-F) Analysis of inherited silencing triggered by λN::HRDE-1 tethering. After outcross to *hrde-1* wild-type (E) or *hrde-1* null (F), reporter worms were scored for *gfp* expression for 13 generations after segregating away the *λN::hrde-1* allele. The percentage of GFP-positive (ON) or GFP-negative (OFF) worms is indicated, N>30 worms scored in each generation. (G) Color chart showing genetic requirements of inherited silencing triggered by λN::HRDE-1 tethering. The *λN::hrde-1; reporter* worms were crossed to the indicated mutants. After segregating away *λN::hrde-1*, reporter worms homozygous for the indicated mutations were scored for GFP expression: ON or OFF, as indicated. N>30 worms scored for each genotype.

Using CRISPR, we inserted an in-frame λN coding sequence at the 5′ end of the endogenous *hrde-1* or *nrde-2* loci (see Methods). Both fusion genes were fully functional, based on their ability to mediate small-RNA guided piRNA and RNAi silencing (Figure S1A). We then tested whether the λN fusions could induce heritable silencing of a reporter gene whose 3′ UTR contains λN-binding sites (i.e., box-B elements) (Figure 1A-D). Both λN::HRDE-1 and λN::NRDE-2 induced silencing of the reporter beginning at the initial heterozygous generation (Figure 1C and 1D). Notably, silencing of the reporter persisted in subsequent generations after genetically segregating away the λN-fusion alleles (Figure 1E, Figure S1B, and data not shown). As expected, inherited silencing (after segregating the λN-fusion alleles) required known components of the transgenerational RNA silencing pathway. For example, HRDE-1 itself was required, as were the small RNA amplification factors RDE-3/MUT-2 and MUT-16 (Gu et al., 2009; Zhang et al., 2011; Shukla et al., 2020) and the nuclear silencing factors NRDE-2 and NRDE-4 (Guang et al., 2010; Ashe et al., 2012) (Figure 1E-G; Figure S1B-G; and data not shown). As expected, λN::HRDE-1 and λN::NRDE-2 tethering induced trimethylation of histone H3 lysine 9 (H3K9me3; Figure 2A-B). Moreover, consistent with the role of H3K9me3 in transcriptional silencing (Buckley et al., 2012), we observed reduced levels of both reporter mRNA and pre-mRNA upon tethering (Figure 2C-D). Thus, artificially recruiting HRDE-1 or NRDE-2 to a target locus was sufficient to initiate the full cycle of events required for inherited silencing, including small RNA amplification and heterochromatin formation.

**Figure 2.**
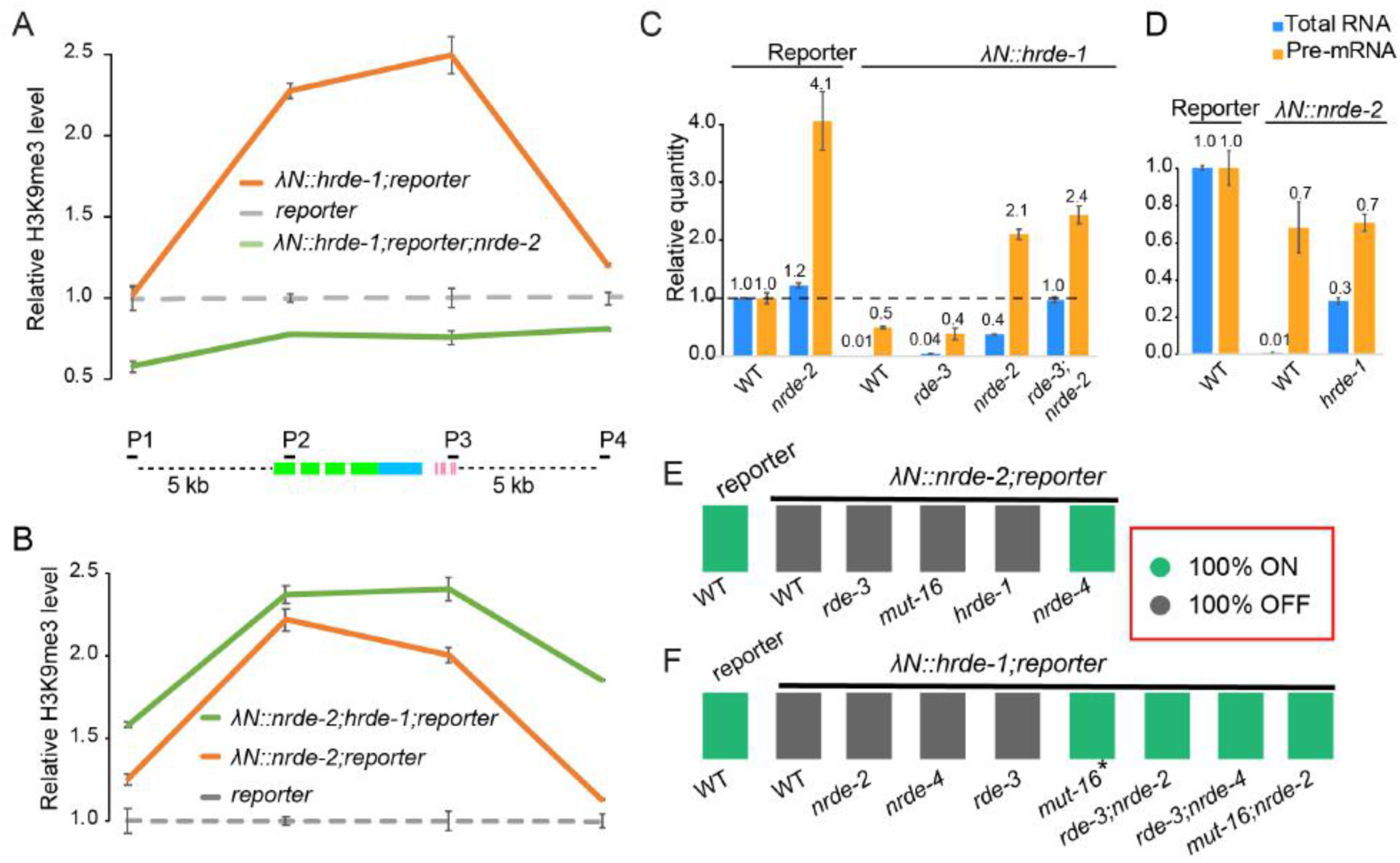
HRDE-1 and NRDE-2 tethering induce heterochromatin formation. (A-B) Quantification of H3K9me3 levels near the reporter in the presence or absence of HRDE-1 or NRDE-2 tethering, as determined by ChIP-qPCR. P1 and P4 primer sets analyze sequences 5 kb upstream or downstream of the reporter, and P2 and P3 analyze sequences within the reporter, as indicated in the schematic. All quantities were normalized to the level of P1 in *reporter* control samples. Error bars show the standard deviation from the mean. (C-D) Bar graphs showing the quantification of reporter RNA and pre-mRNA levels in response to HRDE-1 or NRDE-2 tethering, as determined by qPCR. The average quantities relative to wild type (wt) are indicated. Error bars show the standard deviation from the mean. (E-F) Color chart showing the genetic requirements of silencing in the presence of λN::NRDE-2 or λN::HRDE-1. Reporter worms homozygous for the indicated mutations were scored for GFP expression: ON or OFF, as indicated. N>30 worms scored for each genotype. * GFP is ON, but signal is weak (see Figure S2H).

Having established that tethering induces inherited silencing that depends genetically on known components of the RNA silencing pathway, we asked which factors were required for silencing when the tethered protein was continuously present. For example, because the λN–box-B interaction recruits HRDE-1 and NRDE-2 independently of a guide RNA, we reasoned that the small RNA amplification machinery should be unnecessary when nuclear silencing factors are tethered to the reporter. Consistent with this idea, we found that λN::NRDE-2 silenced the reporter in the absence of *rde-3*, *mut-16*, and *hrde-1* (Figure 2E and Figure S2A-C), but failed to silence in the absence of *nrde-4* (Figure 2E and Figure S2D). These results suggest that NRDE-2 acts downstream of HRDE-1 and upstream of NRDE-4 in nuclear silencing.

Interestingly, silencing by λN::HRDE-1 was maintained independently of *nrde-2*, *nrde*-*4*, or *rde-3*, and only partly required *mut-16* activity (Figure 2F and Figure S2E-H). To completely prevent silencing by λN::HRDE-1, it was necessary to simultaneously mutate components of both the small RNA amplification machinery (*rde-3* or *mut-16*) and components of the chromatin nuclear silencing machinery (*nrde-2* or *nrde-4*) (Figure 2F and Figure S2E-K). Thus, tethering HRDE-1 appears to direct silencing via both the chromatin silencing machinery and the small RNA post-transcriptional silencing pathway (Figure 2A, 2C and 2F).

### HRDE-1 acts downstream of NRDE-2 to promote small-RNA amplification

The above findings suggest that HRDE-1 initiates inherited silencing independently of *nrde-2* and *nrde-4*, while NRDE-2 requires both *nrde-4* and *hrde-1*. A likely explanation for these findings is that heterochromatin silencing directed by NRDE-2 and NRDE-4 induces the de novo synthesis of small RNAs that engage HRDE-1, and that HRDE-1 can further amplify these small RNAs to propagate silencing to offspring. Indeed, whereas we detected very few small RNAs targeting the reporter in the absence of tethering (Figure 3A), tethering NRDE-2 induced small RNA accumulation that required *nrde-4, rde-3*, and *hrde-1* (Figure 3B-E).

**Figure 3.**
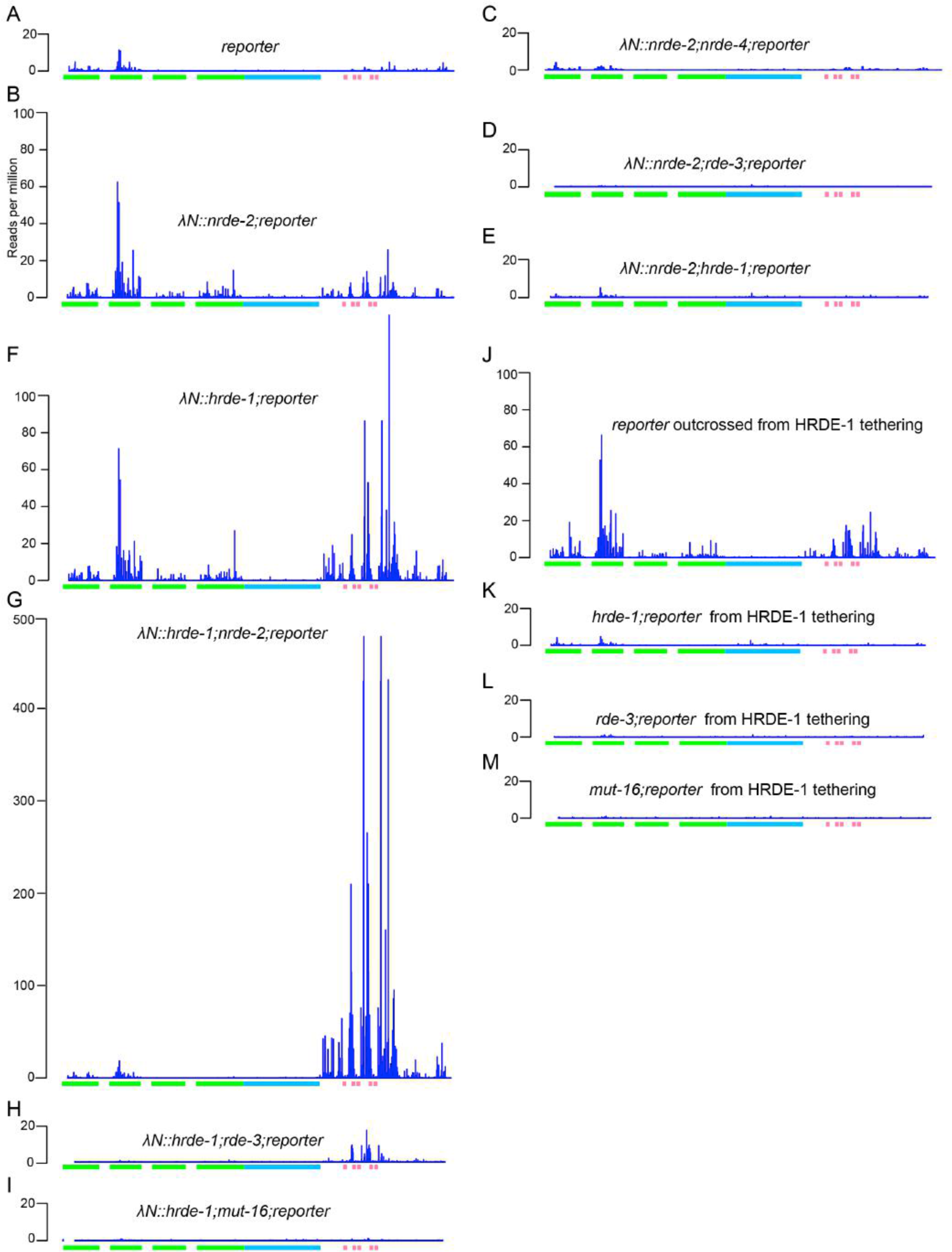
HRDE-1 and NRDE-2 tethering promote antisense small-RNA production. (A) Plot showing antisense small RNA reads (per million total reads) mapping to the reporter (indicated below the plot) in the absence of tethering. Only the 1^st^ nucleotide is counted. Green boxes, GFP coding; blue box, H2B coding; pink boxes, BoxB hairpins. (B-E) Genetic requirements of small RNAs induced by NRDE-2 tethering. Plots showing antisense small RNA reads mapping to the reporter in the presence of λN::NRDE-2 in wild-type (B), *nrde-4* (C), *rde-3* (D), or *hrde-1* (E) worms. (F-I) Genetic requirements of small RNAs induced by HRDE-1 tethering. Plots showing antisense small RNA reads mapping to the reporter in the presence of λN::HRDE-1 in wild-type (F), *nrde-2* (G), *rde-3* (H), or *mut-16* (I) worms. (J-M) Genetic requirements of inherited small RNAs induced by HRDE-1 tethering. Plots showing antisense small RNA reads mapping to the reporter in wild-type (F), *nrde-2* (G), *rde-3* (H), or *mut-16* (I) worms after segregating λN::HRDE-1. Note in (G), y axis is compressed 50% compared to other plots to conserve space.

In contrast, λN::HRDE-1 tethering induced abundant small RNA accumulation that was independent of *nrde-2* and *nrde-4* (Figure 3F-G and S3A). Interestingly, both the distribution of small RNAs and their levels of accumulation along the target mRNA were dramatically altered in the *nrde* mutants after λN::HRDE-1 tethering. Small RNA levels were markedly increased adjacent to the box-B sites and were diminished on the *gfp* coding sequences (Figure 3G and Figure S3B). Small RNAs targeting the reporter were greatly reduced by mutations in *rde-3* and *mut-16*, as expected, (Figure 3H-I). Interestingly, however, a low level of small RNAs persisted directly adjacent to the boxB sites when λN::HRDE-1 was tethered in the absence of *rde-3*, but not in the absence of *mut-16* (Figure 3H). This result is consistent with the observation that tethering of λN::HRDE-1 can bypass *rde-3* but cannot fully bypass *mut-16* (Figure 2F).

When outcrossed to a *hrde-1(+)* background to segregate away λN::HRDE-1, the reporter remained silent for at least 13 generations, with no change in penetrance. Moreover, we observed only a slight reduction in small RNA levels primarily in regions juxtaposed to the box-B hairpins (Figure 3J). In contrast, when outcrossed to a *hrde-1* null background, the reporter was fully de-silenced and small RNAs were absent (Figure 3K). As expected, the maintenance of silencing, and of small RNA levels, also required *rde-3(+)* and *mut-16(+)* (Figure 3L-M). Taken together these findings suggest that chromatin silencing induces de novo transcription and loading of small-RNAs onto the nuclear Argonaute HRDE-1, which further promotes small-RNA amplification (see Discussion).

### HRDE-1 guide-RNA loading is not required for small-RNA amplification

The finding that λN::HRDE-1 can direct chromatin silencing in *rde-3* and *mut-16* mutants, which are defective in small-RNA amplification, suggests that the unloaded Argonaute can direct chromatin silencing when tethered. To further test this idea, we took two approaches. First we monitored silencing in the *hrde-2* mutant (which is defective in HRDE-1 small RNA loading (Spraklin et al., 2017) and second we monitored silencing in λN::HRDE-1(Y669E) a mutant defective in guide RNA binding (Figure S5B) that was modeled on a previous structural study (Rüdel et al., 2011). In both cases, we found that tethering completely silenced the boxB reporter as monitored by GFP fluorescence (Figure 4C and S4D). As expected, the *hrde-1(Y669E)* mutant is defective in silencing a piRNA reporter (Figure S4A) and shows small RNA collapse as in *hrde-1(null)* (Figure S4B and S4C). However, in these mutant contexts, loss of *rde-3* alone was sufficient to completely de-silencing the reporter (Figure S4E and Figure 4C) suggesting that in the absence of guide-RNA loading HRDE-1 fails to engage the NRDE chromatin silencing arm of the pathway. Deep sequencing revealed an abundant accumulation of *rde-3*-dependent small RNAs targeting the boxB reporter in λN::HRDE-1(Y669E) animals (Figure 4E and 4F). Notably, the pattern and levels of small RNA accumulation induced by λN::HRDE-1(Y669E) resembled those observed when wild-type λN::HRDE-1 is tethered in a *nrde-2* mutant (compare Figure 4E to 3G)—i.e., resulting in increased levels of small RNAs targeting sequences adjacent to the boxB sites and reduced levels targeting GFP sequences. Taken together these results suggest that tethering of unloaded HRDE-1 can induce local small RNA amplification and silencing, but that HRDE-1 must be loaded with small RNAs for tethering to induce chromatin silencing and, finally, that chromatin silencing, in turn, is required for small RNA targeting to spread into the 5’ sequences of the target mRNA.

**Figure 4.**
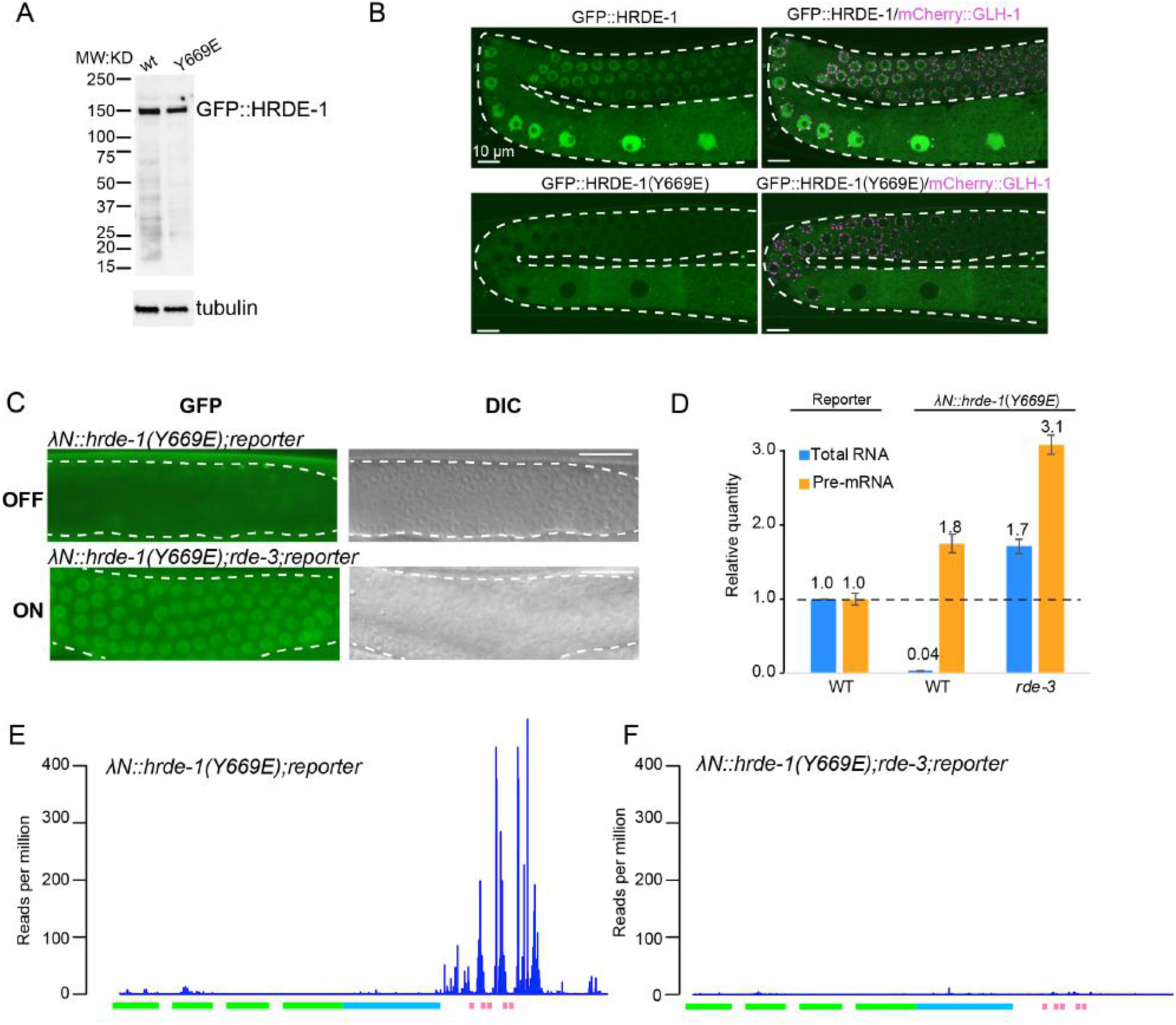
HRDE-1 guide RNA loading is not required for small-RNA amplification. (A) Western blot analysis to detect GFP::HRDE-1 and GFP::HRDE-1(Y669E) in worm lysates. Upper panel probed with anti-GFP antibody. GFP::HRDE-1 was indicated. Lower panel probed with anti-Tubulin antibody as a loading control. (B) Confocal images showing the localization of GFP::HRDE-1(wild-type) or GFP::HRDE-1(Y669E) with mCherry::GLH-1 as P-granule marker. The white dashed lines outline a gonadal arm of the germline. (C) Representative fluorescence (left panels) and DIC (right) images showing that λN::HRDE-1(Y669E) silences the BoxB reporter in wild-type worms (upper panels, OFF), but not in *rde-3* mutant worms (lower panels, ON). The images represent 100% of the animals scored, N>30. (D) Bar graphs showing the quantification of reporter RNA and pre-mRNA levels in response to HRDE-1(Y669E) tethering, determined by qPCR. The average quantities relative to wild type (wt) are indicated. Error bars show the standard deviation from the mean. (E and F) Plots showing antisense small RNA reads mapping to the reporter in the presence of λN::HRDE-1(Y669E) in wild-type (E) or *rde-3* (F) worms.

### HRDE-1 promotes small RNA amplification through its N-terminal domain

We next attempted to dissect functional domains of HRDE-1 required for small RNA amplification. We used CRISPR to make a series of *λN::hrde-1* truncation mutants (Figure 5A). These studies identified the N-terminal half (herein N-terminal domain, NTD) as the minimal fragment of HRDE-1 that could fully silence the reporter. The NTD and the remaining C-terminal domain (CTD) truncations of HRDE-1 are predicted by I-TASSER (Yang and Zhang, 2015) to fold into self-contained globular structures, with subdomains similar to those identified in atomic resolution studies on Human Ago2 (Schirle et al., 2014) (Figure 5B and Figure S5A-B). As expected, in the absence of tethering, *hrde-1(NTD)* and *hrde-1(CTD)* alleles failed to silence a piRNA sensor (Figure S4A).

**Figure 5.**
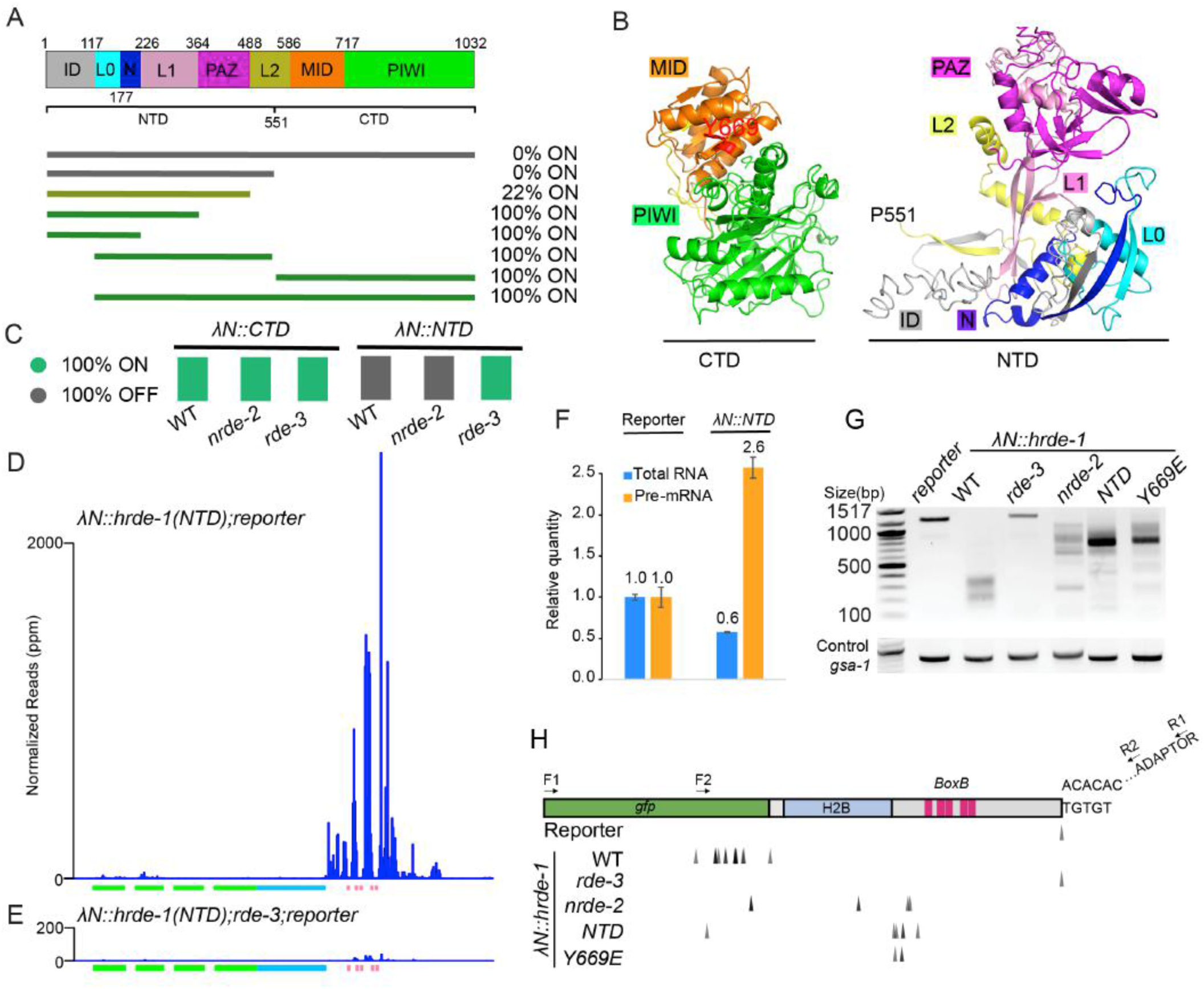
HRDE-1 N-terminal domain promotes small-RNA amplification and poly-UG modification. (A) Schematic showing HRDE-1 linear domain structure and truncations tested. The subdomains are color-coded based on human Ago2 (Figure S5A). The percentage of GFP-positive worms (ON) is indicated, N>30 worms scored in each test. (B) Predicted three-dimensional structures of HRDE-1 N-terminal domain (NTD) and C-terminal domain (CTD). Subdomains as in Figure 5A. (C) Color chart indicating the expression (ON) or silencing (OFF) of the reporter in the presence of λN::CTD or λN::NTD and the requirement of *nrde-2* or *rde-3*. N>30 worms scored for each genotype. (D-E) Plots showing antisense small RNA reads mapping to the reporter in the presence of λN::NTD in wild-type (D) or *rde-3* worms (E). (F) Bar graphs showing the quantification of reporter RNA and pre-mRNA levels in response to NTD tethering, as determined by qPCR. The average quantities relative to the control are indicated. Error bars show the standard deviation from the mean. (G-H) Analysis of poly-UG modification of reporter RNA in response to tethering in the indicated mutants. Poly-UG PCR products in (G) were cloned and sequenced to identify the precise positions of poly-UG addition (H), indicated by arrowheads. A *gsa-1*-specific PCR was used as loading control.

Silencing by λN::NTD required *rde-3* but not *nrde-2* (Figure 5C and Figure S5C), and deep sequencing revealed that λN::NTD induces abundant *rde-3-*dependent small RNAs targeting the box-B reporter (Figure 5D and 5E). Truncations that failed to silence the reporter did not trigger small RNA generation (Figure S5D). The small RNA pattern induced by λN::NTD resembled the patterns caused by λN::HRDE-1 in *nrde-2* mutants or by λN::HRDE-1(Y669E)—i.e., dramatically increased level of small RNAs proximal to the boxB sites and reduced levels of small RNAs targeting GFP sequences. Interestingly, the magnitude of small RNA accumulation induced by λN::NTD at the boxB sites was ~4-fold greater than that induced by either λN::HRDE-1 in *nrde-2* mutants or by λN::HRDE-1(Y669E) (compare Figure 5D to Figures 3G and 4E). These results suggest that the NTD of HRDE-1 robustly recruits the small RNA amplification machinery to the target and promotes silencing that is independent of the nuclear silencing pathway.

### HRDE-1 tethering promotes accumulation of poly-UG-modified target fragments

During RNA silencing in worms, truncated target RNAs are converted into templates for small RNA production via the RDE-3-dependent addition of poly-UG tails (Shukla et al., 2020). We therefore used a quantitative PCR assay (Shukla et al., 2020) to detect poly-UG additions to reporter RNA in the absence of a λN fusion or in worms expressing λN::HRDE-1, λN::NTD, or λN::HRDE-1(Y669E) (Figure 5G and 5H). Priming from an endogenous UGUG motif in the reporter 3′ UTR serves as a control for the presence of full-length mRNA. This analysis revealed that faster-migrating, poly-UG-modified RNAs accumulated in strains where silencing was active. In wild-type λN::HRDE-1 worms, poly-UG-modified RNAs were most robustly detected at truncations within the GFP sequences (Figure5G and 5H). As expected, only full-length mRNA was detected in *rde-3* mutants, confirming that RDE-3 is absolutely required for poly-UG RNA accumulation. Notably, mutation of *nrde-2* or tethering the NTD or Y669E mutants shifted poly-UG addition toward the 3′ end of the reporter, close to the boxB elements (Figure 5G and 5H). These results suggest that HRDE-1 tethering induces RDE-3-dependent poly-UG modification of truncation products that are generated near the tethering sites and that nuclear silencing promotes the induction of additional truncations far away from the tethering sites that likely support the 5’ spread of small RNA amplification.

To further analyze changes in target RNA caused by tethering we used real-time quantitative (RT-qPCR) and Northern blot analysis. As expected, tethering wild-type λN::HRDE-1 dramatically reduced the reporter mRNA by 99% as measured by RT-qPCR, while pre-mRNA levels were reduced by 50% (Figure 2C). Surprisingly, λN::NTD actually caused an approximately 2.5-fold increase in pre-mRNA levels and only a 40% decrease in total level, as judged by RT-qPCR (Figure 5F). The latter result was surprising given that GFP fluorescence was indetectable in λN::NTD worms (Figure 5C and Figure S5C) and suggested that the accumulating species in λN::NTD animals might reflect the accumulation of the nearly full-length pUG RNA. Consistent with this idea, Northern blot analysis detected a reduced level of an slightly shorter reporter mRNA species in the λN::NTD animals (Figure S5E).

### Functional HRDE-1 RISC is not required parentally for transmission of silencing to offspring

We next wished to ask if λN::NTD can initiate inherited silencing. To do this we first established reporter silencing by tethering λN::NTD in an otherwise wildtype background. We then crossed to a reporter strain homozygous for a *hrde-1* null allele to generate animals heterozygous for the tethering construct. Finally, we crossed these heterozygotes (either as males or hermaphrodites) to a reporter strain wild-type for *hrde-1(+)*. These steps generated two types of cross progeny; λN::NTD/+ or null/+ heterozygotes, that were derived from parents baring a silent reporter but lacking a copy of HRDE-1 capable of binding to guide RNA and hence of assembling a functional RNA-induced Silencing Complex (RISC). We found that male and female gametes that lacked a functional HRDE-1 RISC nevertheless robustly transmitted silencing to the next generation (Figure S7A and S7B). As expected HRDE-1(+) was required in the inheriting generation for silencing to occur ((Buckley et al., 2012) and Figure1F). Since the NTD fails to establish heterochromatin upon tethering and cannot directly form a RISC complex, these findings suggest that parentally established heterochromatin and HRDE-1 RISC are not required in gametes for inheritance, a finding consistent with previous work in which *hrde-1(−/−)* hermaphrodites were shown to transmit silencing to their *hrde-1(+/−)* progeny (Buckley et al). Rather, in the parental generation the tethered NTD can stimulate amplification of small RNAs that likely engage with other Argonautes, to propagate silencing to offspring (see Discussion).

### HRDE-1 localizes to Mutator foci

HRDE-1 localization is primarily nuclear (Buckley et al., 2012), however, template formation and small-RNA amplification are thought to occur in Mutator foci, domains of peri-nuclear nuage where several components of the small RNA-amplification machinery localize (Zhang et al., 2011; Phillips et al., 2012; Shukla et al., 2020). To examine whether HRDE-1 might also localize in Mutator foci, we expressed GFP::HRDE-1 (without tethering) in worms that also express either mCherry::GLH-1, which localizes broadly within nuage, or MUT-16::mCherry, which localizes prominently in Mutator foci. GFP::HRDE-1 co-localized to a subset of peri-nuclear mCherry::GLH-1 foci, especially in association with late pachytene germ nuclei (Figure 6A and Figure S6A). Moreover, the GFP::HRDE-1 foci only partially overlapped with mCherry::GLH-1 foci, suggesting that the HRDE-1-positive foci occupy subdomains of larger GLH-1-positive nuage, reminiscent of Mutator foci. Indeed, GFP::HRDE-1 foci coincided almost perfectly with MUT-16::mCherry foci (Figure 6B). Similarly, GFP::HRDE-1(NTD) colocalized with GLH-1::mCherry and mCherry::MUT-16 foci (Figure 6C-D and FigureS6B). Taken together, these findings suggest that HRDE-1 localizes via its NTD to Mutator foci where it functions to promote small-RNA amplification.

**Figure 6.**
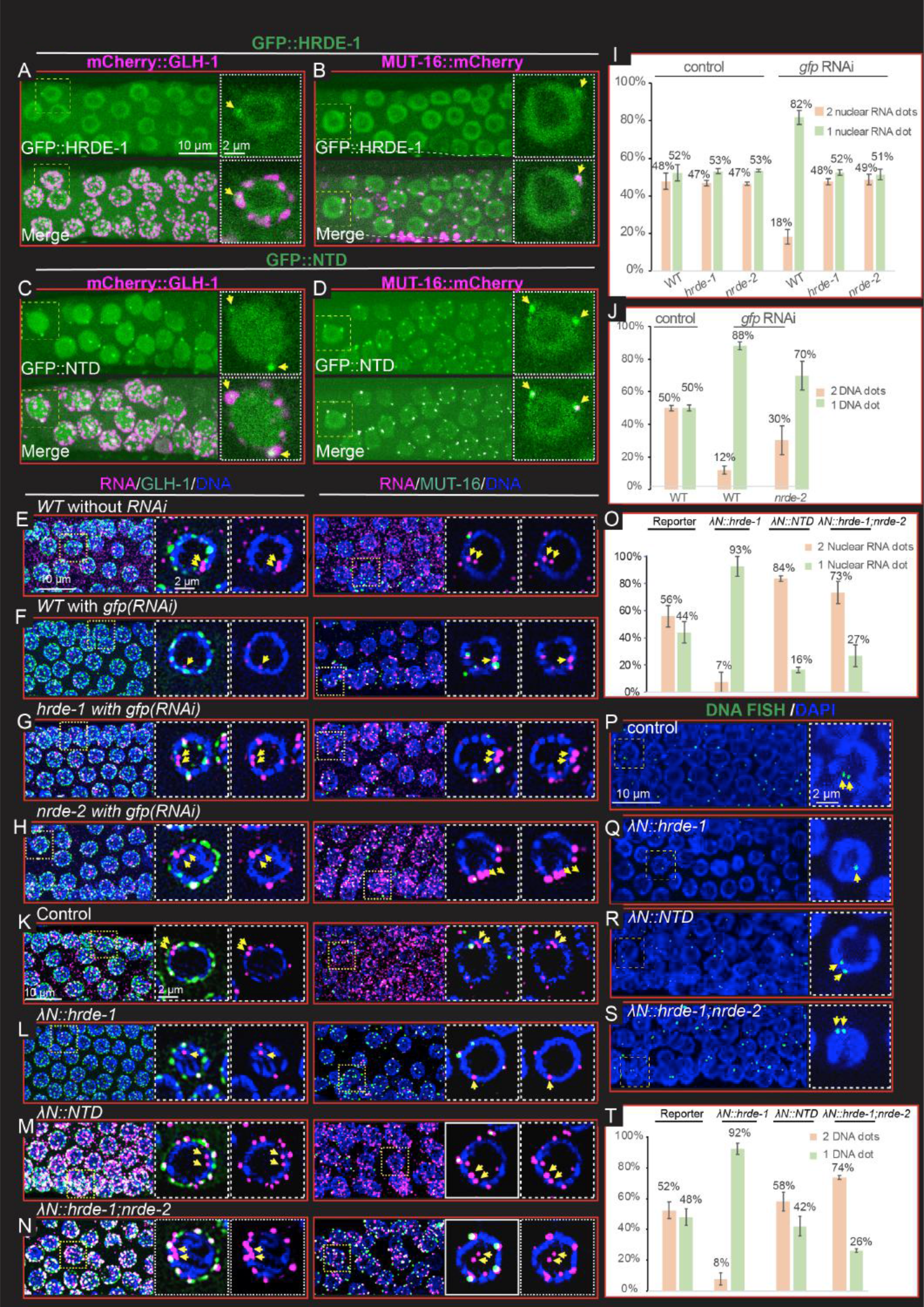
HRDE-1 localizes in Mutator foci and HRDE-1 tethering caused peri-nuclear accumulation of reporter RNA in nuclear silencing mutants. (A-B) Confocal image of live germ cells showing the co-localization of GFP::HRDE-1 with mCherry::GLH-1 (A) and MUT-16::mCherry (B). Each subpanel shows a projected view of a segment of the germline to the left, and the nucleus bounded by a dashed box is shown as a single-focal-plane image to the right. Yellow arrows point to perinuclear foci where HRDE-1 co-localizes with GLH-1 and MUT-16. (C-D) Confocal images of live germ cells showing the co-localization of GFP::HRDE-1(NTD) with mCherry::GLH-1 (C) and MUT-16::mCherry (D). As in panels A and B. (E-F) Confocal images of RNA FISH experiments showing the localization of reporter RNA with mCherry::GLH-1 (left) or MUT-16::mCherry (right) in control worms (E) or in worms exposed to *gfp* RNAi (F). Magenta, RNA; green, GLH-1 or MUT-16; and blue, DAPI. Each subpanel shows a projected view of a segment of a representative germline to the left, and the nucleus bounded by a dashed box is shown as a single focal-plane image with DNA and GLH-1 or MUT-16 signals (center) or with DNA signal only (right). Yellow arrows point to nuclear RNA foci that likely correspond to transcription sites. (G-H) As in panel F, but in *hrde-1* (G) or *nrde-2* (H) mutant worms. (I) Bar graphs showing the percentage of nuclei from panels E to H containing one reporter RNA focus (orange) or two or more reporter RNA foci (light green). Three independent germlines were measured for each condition. (J) Bar graphs showing the percentage of nuclei from DNA FISH (Figure S6E) containing one reporter DNA focus or two or more reporter DNA foci. Similar to panel I. (K-N) Confocal images of RNA FISH showing the localization of reporter RNA with mCherry::GLH-1 (left) or MUT-16::mCherry (right) in the absence (K) or presence (L-N) of HRDE-1 tethering, as indicated. Details as in panels E-F. (O) Bar graphs showing the percentage of nuclei from panels J to N containing one reporter RNA focus (peach) or two or more reporter RNA foci (light green). (P-S) Confocal images of DNA FISH experiments showing the localization of reporter DNA loci in the absence (P) or presence (Q-S) of HRDE-1 tethering, as indicated. Green, DNA FISH signal; blue, DAPI. A projected view of a segment of a representative germline is shown to the left, and the nucleus bounded by a dashed box is shown as a single-focal-plane image to the right. Yellow arrows point to the nuclear DNA signals. (T) Bar graphs showing the percentage of nuclei from DNA FISH experiments in panels P to S containing one reporter DNA focus or two or more reporter DNA foci. Details as in panel I.

### Silencing by dsRNA or tethering causes target genes to co-localize

To understand how HRDE-1 and nuclear silencing regulate their target genes and RNAs, we performed RNA and DNA fluorescence in situ hybridization (FISH) studies to visualize the boxB reporter mRNA and DNA. In the absence of silencing, reporter RNA foci were detected throughout the germline cytoplasm (Figure 6E and Figure S6C). In addition, we observed prominent RNA signals in the majority (~70%) of pachytene nuclei (most nuclei, 57%, exhibited at least two closely paired nuclear dots, while the remainder exhibited a single dot; Figure 6E and 6I). The positions of these nuclear signals adjacent to DAPI-stained chromosomes suggests that they correspond to sites of transcription on the paired sister chromatids within the axial loops of synapsed meiotic homologs. Silencing induced either by exposure to dsRNA targeting the reporter or by tethering λN::HRDE-1 eliminated cytoplasmic reporter RNA signal and greatly reduced the nuclear signal (Figure 6F, 6L and S6C). More than 80% of the pachytene nuclei with visible RNA signal exhibited a single nuclear focus (Figure 6F, 6L, 6I and 6O). The changes in nuclear RNA signal induced by silencing correlated with changes in the reporter DNA FISH signal. In the absence of silencing, we observed a pair of nuclear DNA FISH signals in approximately 50% of pachytene nuclei that have visible DNA signal (Figures 6P and Figure 6T), while in the presence of silencing, we observed a single focus of DNA FISH signal in approximately 90% of pachytene nuclei with visible DNA signal (Figure 6Q, 6J, 6T and S6E). These results suggest that nuclear silencing mediated by HRDE-1 causes the target alleles, which consist of 4 genes resolved prior to silencing as two nearby DNA FISH signals (each signal comprised of two closely localized sister chromatids), merge from these predominantly paired DNA fish signals into a single focus containing all 4 silenced alleles.

### Mutations that disarm nuclear silencing cause target RNA to accumulate in nuage subdomains that resemble Mutator Foci

We next examined how mutations that disarm only the nuclear silencing pathway impact RNA and DNA localization after RNAi or tethering. To do this we performed RNA and DNA FISH on λN::NTD worms and on *nrde-2* mutants. In these mutants where nuclear silencing is disarmed, we found that nuclear RNA and DNA FISH signals resembled the nuclear signals observed in wild-type animals in the absence of silencing: exhibiting predominantly two foci of RNA and DNA FISH signals in each background (Figure 6M, N, I, J, O, T and S6D). Strikingly, however, while RNA signal was absent from the bulk cytoplasm throughout the gonad, consistent with cytoplasmic post-transcriptional silencing, we noticed pronounced accumulation of reporter RNA signals in multiple peri-nuclear foci surrounding pachytene nuclei. Co-staining experiments with GFP::GLH-1 or MUT-16::GFP revealed that these RNA foci coincide with most of the nuage sub-domains that express MUT-16::GFP (Figure 6G-H and 6M-N). The accumulation of target RNA in the MUT-16 foci required RDE-3 (Figure S6F), suggesting that these RNA signals may correspond to RdRP templates engaged in small-RNA amplification.

### MUT-16 promotes the nuclear localization of GFP::HRDE-1 but not its nuage localization

MUT-16 is required for the co-localization of small-RNA amplification factors within Mutator foci (Zhang et al., 2011; Phillips et al., 2012; Uebel et al., 2018). We therefore wondered if MUT-16 is also required for the co-localization of HRDE-1 in Mutator foci. To answer this question we introduced a null allele of *mut-16* into worms expressing both GFP::HRDE-1 and mCherry::GLH-1. As shown previously (Cornes et al., 2022), we found that MUT-16 activity is required for the nuclear localization of HRDE-1 (Figure 7A-B). MUT-16 was not however required for the localization of GFP::HRDE-1 to nuage (Figure 7A-B). The localization of GFP::HRDE-1 in nuage appeared more obvious in *mut-16* mutants, but the levels of GFP::HRDE-1 within nuage and the approximate numbers of foci appeared similar with or without *mut-16* activity (Figure 7A-B). Finally, the localization of MUT-16 itself to nuage was not disrupted in *hrde-1* mutants (data not shown), thus HRDE-1 and MUT-16 localize within a nuage subdomain (or domains) independently of each other.

**Figure 7.**
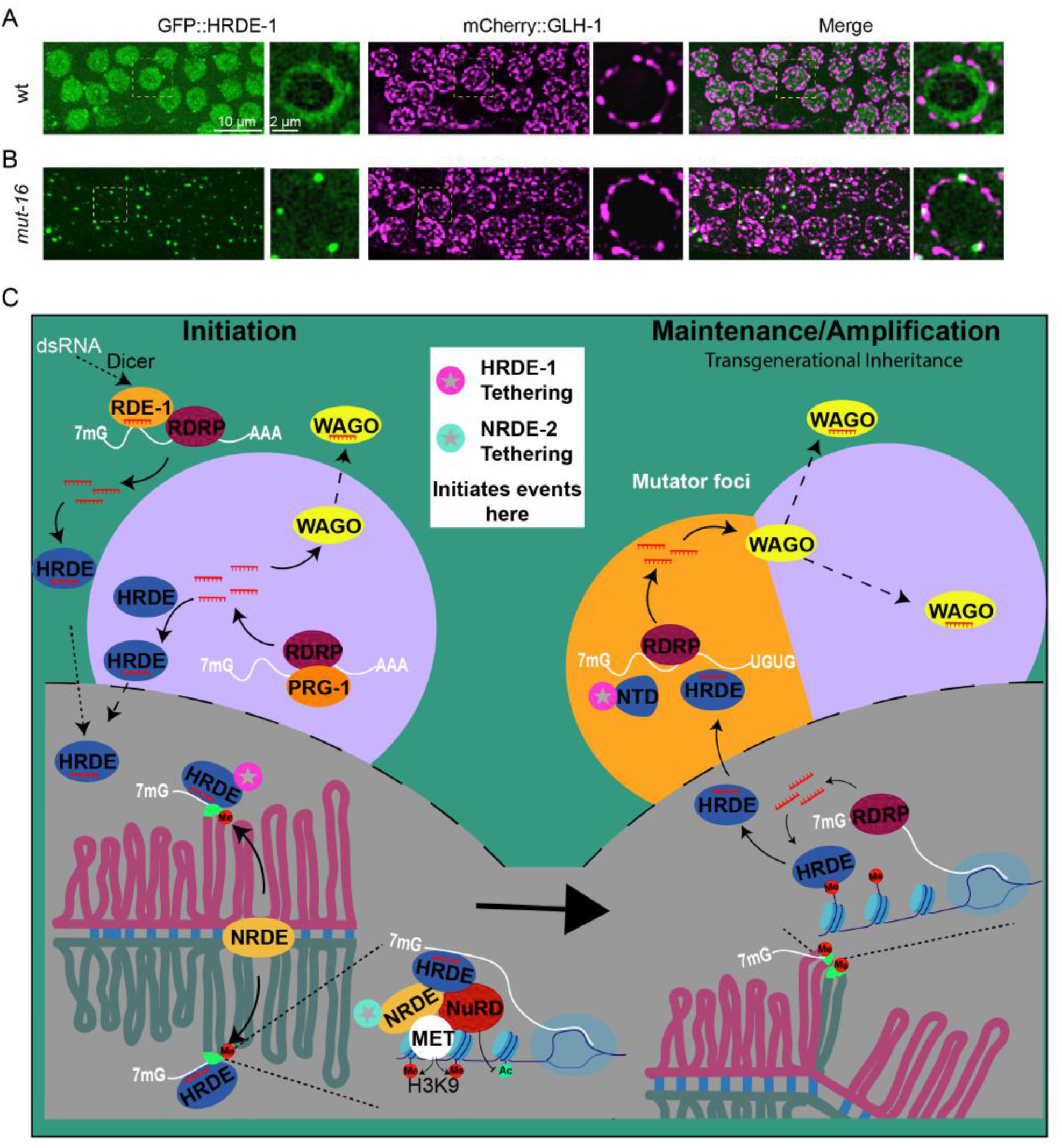
Model of HRDE-1 mediated self-enforcing mechanism. (A-B) Confocal images showing the localization of GFP::HRDE-1 with mCherry::GLH-1 in wild-type worms (A) and *mut-16* mutants (B). Green, GFP::HRDE-1 (left); magenta, mCherry::GLH-1 (middle); merge (right). Each subpanel shows a projected image of a representative pachytene region of the germline to the left, and the nucleus bounded by a dashed box is shown as a single-focal-plane image to the right. (C) Model (see Discussion).

## Discussion

In many eukaryotes the installation and maintenance of chromatin silencing is coupled to Argonaute small-RNA pathways that promote transmission to offspring. Here we have explored the role of a nuclear Argonaute HRDE-1 in coordinating transgenerational silencing in the *C. elegans* germline. In addition to its known role in directing heterochromatin silencing downstream of RNAi (Buckley et al., 2012; Spracklin et al., 2017) and Piwi Argonaute silencing (Ashe et al., 2012; Shirayama et al., 2012; Sapetschnig et al., 2015), our tethering studies have shown that HRDE-1 is also loaded with small-RNA de-novo, downstream of heterochromatin silencing enabling it to prime a new round of small-RNA amplification within nuage (Figure 7C:Model).

The nuclear silencing events that depend on HRDE-1 cause the target alleles to co-localize into a single focus of DNA FISH signal (Figure 6P-S and Figure S6E). Presumably, the heterochromatinized alleles within this focus are transcribed at low levels to produce template RNA that feeds transgenerational silencing (Holoch and Moazed, 2015). Consistent with this idea, the inactivation of nuclear silencing caused target alleles to remain separated and increased the levels of the nuclear- and nuage-localized RNA signals. The failure to engage nuclear silencing and the consequent increased expression of RNA that feeds template formation in nuage likely explains why small-RNA levels are dramatically increased when HRDE-1-dependent nuclear silencing activities are disarmed by mutation.

In the yeast *S. pombe*, the RNAi-induced transcriptional silencing complex (RITS), which includes an RdRP and a nuclear Argonaute AGO1p, resides in heterochromatin. Tethering AGO1p to RNA using a box-B reporter system, similar to the one used here, was sufficient to recruit the RITS complex, induce small RNA amplification, and to drive reporter silencing (Bühler et al., 2006).

HRDE-1 associates with NRDE-2 and components of the Nucleosome Remodeling and Deacetylase NuRD complex to establish heterochromatin silencing (Buckley et al., 2012; Wan et al., 2020; Kim et al., 2021). However, how heterochromatin, once formed, leads to de-novo programming of HRDE-1 is unknown. In *C. elegans*, the RdRP EGO-1 has been shown to associate with germline chromatin (Maine et al., 2005; Claycomb et al., 2009), and several of our findings would be consistent with a cycle of nuclear small-RNA transcription and de novo HRDE-1 loading within heterochromatin. Such a mechanism could explain why tethering NRDE-2 in the absence of HRDE-1 initiates heterochromatin silencing but not small RNA amplification (Figure 2E and Figure 3E). Perhaps after a nuclear cycle of HRDE-1 loading the protein exits the nucleus along with nascent target/template RNA to further amplify small-RNA production. Consistent with this idea, we have shown that the N-terminal half of HRDE-1 (its NTD) is sufficient to stimulate small-RNA amplification and loading, and that both the NTD and full-length HRDE-1 (as well as target RNA) localize within nuage, within a specialized nuage domain known as Mutator foci.

Mutator foci accumulate poly-UG modified templates derived from target RNA (Shukla et al., 2020) and are thought to serve in the amplification of small-RNA signals that are propagated to offspring. Thus, our findings suggest that HRDE-1 shuttles out of the nucleus to nuage to promote small-RNA amplification. A mutant HRDE-1 protein incapable of binding guide RNA was sufficient (when tethered) to induce silencing that transmits to offspring via either the sperm or the egg (Figure S7A-B), thus, as previously reported (Buckley et al., 2012) a functional HRDE-1 RISC is not required in gametes for transgenerational silencing. Instead, functional HRDE-1 capable of binding guide RNA was required in offspring to renew silencing for another generation (Buckley et al., and Figure 1F). Thus, in the parent, Mutator foci likely serve as locations where HRDE-1 and other upstream Argonautes trigger the expansion of small RNAs that are loaded onto downstream WAGO Argonautes, including the two prominent nuage localized WAGOs WAGO-1 (Shirayama et al., 2012) and WAGO-4 (Xu et al., 2018). Consistent with this idea, silencing induced by λN::HRDE-1(Y669E) was partially dependent on *wago-1* (75% de-silenced, N=32 and Figure S4G).

Taken together our findings suggest that heterochromatin renews small-RNA silencing (and vice versa) during each germline life cycle. For example, small RNAs guide heterochromatin formation in the zygote, and heterochromatin then propagates silencing before feeding back into the de-novo synthesis of guide RNAs that load onto HRDE-1. HRDE-1 promotes expansion of small RNAs that are then transmitted to offspring through HRDE-1 and other WAGOs to re-establish heterochromatin. Heterochromatin then, in turn, transcribes RNA that forms templates for RdRP-dependent amplification, renewing the cycle. Consistent with these ideas, neither pathway, small RNA or heterochromatin alone, is sufficient to stably transmit silencing signals for multiple generations (Ashe et al., 2012; Buckley et al., 2012; Shirayama et al., 2012; Spracklin et al., 2017) (Figure S7C-F). Given the similarities between the worm and yeast mechanisms, and by extension the intriguing relationships between long-non coding RNAs and chromatin modifiers in fly and mammal (Weick and Miska, 2014), it seems likely that feedforward RNA-chromatin circuits that amplify and maintain silencing across cell divisions, or generations, will be a common feature of gene regulation in eukaryotes.

## KEY RESOURCE TABLE

| REAGENT or RESOURCE | SOURCE | IDENTIFIER |
| --- | --- | --- |
| Antibodies |  |  |
| Anti-GFP | Cell Signaling | Cat # 2956S |
| Anti-Tubulin | Abcam | Cat # ab6160 |
| Anti-H3K9me3 | Upstate | Cat# 07523 |
| Bacterial Strains |  |  |
| *E. coli*: Strain OP50 | *Caenorhabditis*<br>Genetics Center | WormBase:<br>OP50 |
| *E. coli*: Strain HT115 | *Caenorhabditis*<br>Genetics Center | WormBase:<br>HT115 |
| **Chemicals, Peptides, and Recombinant Proteins** |  |  |
| Alt-R® S.p. Cas9 Nuclease V3 | Integrated DNA<br>Technologies<br>(IDT) | Cat# 1081059 |
| CRISPR-Cas9 tracrRNA | IDT | Cat# 1072534 |
| Tetramisole hydrochloride | Sigma-Aldrich | Cat# L9756 |
| TRI reagent (TRIZOL) | Sigma-Aldrich | Cat# T9424 |
| DNase I | NEB | Cat# M0303S |
| SuperScript™ IV Reverse Transcriptase | Life Technologies | Cat# 18091200 |
| Fast SYBR Green Master Mix | Life Technologies | Cat# 26616 |
| Pierce CHIP grade IgA/G magnetic beads | Life Technologies | Cat# 26162 |
| RNase A | Life Technologies | Cat# AM2270 |
| Protase K | Sigma Aldrich | Cat# RPROTK-RO |
| SUPERase RNase Inhibitor | Invitrogen | Cat# am2696 |
| mirVana™ miRNA Isolation Kit | Life Technologies | Cat# AM1561 |
| T4 RNA Ligase 2, truncated KQ | NEB | Cat# M0373L |
| T4 RNA Ligase 1 | NEB | Cat# M0204S |
| SuperScript™ III Reverse Transcriptase | Life Technologies | Cat# 18080085 |
| NextSeq 500/550 High Output Kit v2.5 (75 Cycles) | Illumina, Inc. | Cat# 20024906 |
| Epredia™ Rite-On Frosted Thick Slide | Fisher Scientific | Cat# 1256820 |
| TOPO™ TA Cloning™ Kit | Life Technologies | Cat# 450641 |
| Taq DNA polymerase | NEB | Cat# M0273S |
| Isopropyl-β-D-thiogalactoside (IPTG) | Sigma-Aldrich | Cat# 11411446001 |
| Tween 20 | Fisher Scientific | BP337-500 |
| Triton X-100 | Sigma-Aldrich | Cat# T8787-250ml |
| KPL Detector™ AP Chemiluminescent Blotting Kit | SeraCare | Cat# 5910-0029 |
| Saline-Sodium Citrate (SSC), 20X Solution | Fisher Scientific | Cat# BP1325-1 |
| Formaldehyde | Sigma-Aldrich | Cat# 252549 |
| Stellaris® RNA FISH Hybridization Buffer | LGC Biosearch, Genomics Ltd | Cat# SMF-HB1-10 |
| VECTASHIELD® Antifade Mounting Medium with DAPI | Vector Laboratories | Cat# H-1200-10 |
| **Deposited Data** |  |  |
| Small RNA sequencing data | This study | NCBI (PRJNA874806) |
| **Strains** |  |  |
| List of *C. elegans* strains | This study | Table S1 |
| **Oligonucleotides** |  |  |
| List of Oligonucleotides | This study | Table S2 |
| List of RNA FISH probes | This study | Table S3 |
| List of DNA FISH probes | This study | Table S4 |
| **Software and Algorithms** |  |  |
| ZEN2 pro | ZEISS | Version blue edition |
| Fusion | Oxford Instruments | Version 2.3.0.44 |
| Microscopy Image Analysis Software: Imaris | Oxford Instruments | Version 9.7.1 |
| Cutadapt | (Magoč and Salzberg, 2011) | Version 4.1 |
| Bowtie2 | (Langmead and Salzberg, 2012) | Version 2.4.5 |
| R | (Team, 2013) | Version 4.2.1 |
| R studio | (Team, 2021) | Version Build 554 |
| DESeq2 | (Love et al., 2017) | version 1.26.0 |
| I-TASSER | (Yang and Zhang, 2015) |  |
| PyMOL(TM) Molecular Graphics System | (Schrodinger, 2010) | Version 2.5.0 |

## METHOD DETAILS

### C. elegans strains and genetics

All the strains used in this study were derived from Bristol N2 and cultured on nematode growth media (NGM) plates with OP50 at room temperature unless indicated otherwise, Strains used in this study were listed in Table S1.

### CRISPR/Cas9 genome editing

The Cas9 ribonucleoprotein (RNP) CRISPR strategy (Dokshin et al., 2018) were used to edit the genome. Plasmid pRF4 containing *rol-6 (su-1006)* was used as co-injection marker. For short insertions like λN and deletion mutations, synthesized single strand DNAs were used as the donor; for long insertions like GFP, mCherry, and 5xBoxB, the annealed PCR products were used instead. The gRNA and donor sequences were listed in Table S2. The BoxB reporter strain was constructed based on a single copy insertion of *Ppie-1:GFP::his-58:unc-54UTR* (WM701). The 5xBoxB sequence amplified from a previously published strain JMC002 (Wedeles et al., 2013) was inserted before the *unc-54* UTR.

### Live Worm fluorescent image

Young adult worms were transferred to glass slide in M9 buffer with 0.4mM Tetramisole. Epifluorescence and differential interference contrast (DIC) microscopy were performed on a Zeiss Axio Imager M2 Microscope and images were processed with ZEN Microscopy Software (Zeiss). Confocal images were taken by a Andor Dragonfly Spinning Disk confocal microscope. Confocal images were processed with Imaris Microscopy Image Analysis Software.

### Quantifying reporter RNA using qPCR

Young adult worms were collected and washed with M9 for three times and ddH_2_O once. Total RNA was extracted with TRIZOL and treated with DNase I to remove DNA contamination. First strand cDNA was synthesized by Superscript IV with random hexamers. Quantitative PCR was performed on a Quant studio 5 Realtime PCR machine together with Fast SYBR Green Master Mix. Actin was used as internal reference (primer set S5265 and S527). Primer set of oYD826 and oYD827 were used for reporter. All primers used were listed in Table S2.

### CHIP-qPCR

A traditional worm CHIP method (Askjaer et al., 2014) was applied to the young adult worm samples. Anti H3K9me3 antibody (Upstate 07523) and CHIP grade IgA/G magnetic beads were used for the immunoprecipitation. During elution, RNase A and Protease K were used to remove RNA and proteins. For qPCR, actin was used as internal reference. All primers used were listed in Table S2.

### Small RNA cloning and data analysis

Small RNA cloning was conducted as previously reported (Kim et al., 2021). Synchronized young adult worms were collected and total RNA were purified with Trizol. Small RNAs were enriched using a mirVana miRNA isolation kit. Homemade PIR-1 was used to remove the di or triphosphate at the 5’ to generate 5’ monophosphorylated small RNA. Adaptors of 3’ (/5rApp/TGGAATTCTCGGGTGCCAAGG/3ddC/) and 5’ (rGrUrUrCrArGrArGrUrUrCrUrArCrArGrUrCrCrGrArCrGrArUrCrNrNrNrCrGrArNrNrNrUr ArCrNrNrN, N is for a random nucleotide) were ligated to the small RNA by T4 RNA ligase 2 (NEB) and T4 ligase 1 (NEB) sequentially. Reverse transcription was performed with SuperScript III and RT primer (CCTTGGCACCCGAGAATTCCA). After PCR amplification, productions around 150 bp were separated by 12% SDS-PAGE and equally mixed. Libraries were sequenced on a NextSeq 550 sequencer with the illumina NextSeq 500/550 high output kit in 75bp single-end sequencing mod. Reads were trimmed by cutadapt and mapped using Bowtie2. For small RNAs mapped to the reporter, total reads with length longer than 16 nt were used to normalized between samples. Plots were generated by R and R studio. DEseq2 package in R was used to find the genes with increased or decreased antisense small RNAs with a criterion of 2-fold change and P value less than 0.05.

### Structure prediction

The 3D structure of HRDE-1 was predicted by I-TASSER online server (Yang and Zhang, 2015) with default setting. HRDE-1 structure was aligned with hAgo2 by PyMOL (Schrodinger, 2010) and its domains were annotated based on the alignment.

### pUG RNA analysis

As previously reported(Shukla et al., 2020), total RNAs were extracted with Trizol. SuperScript IV was used to generate the first strand DNA with reverse transcription primer oYD1001. A pair of outer primers (oYD998 and oYD1002) were used for the first round PCR amplification with Taq DNA polymerase. After 100-fold dilution, another round of PCR was performed with a pair of inner primers (oYD256 and oYD1003). PCR products were analyzed by 1.5% agarose gels. DNA bands were purified, cloned with TOPO TA Cloning Kit and sent for sanger sequencing. *gsa-1* served as a control for pUG PCR analysis.

### Northern blot

Norther blot was done with KPL AP Chemiluminescent Blotting Kit following its instructions. Total RNA of 20 ug was ran in a 1% agarose gel and transferred to positively charged nylon membrane. The RNA was hybridized with homemade biotin labeled anti-*gfp* probe generated from reverse transcription. Alkaline phosphatase-labeled streptavidin boud to the probe and generated signal when incubated with chemiluminescent substrate. Signal was captured with a ChemiDoc gel imaging system (Bio-Rad).

### RNA FISH

Worms at young adult stage were dissected in Happy Buffer (81mM HEPES pH 6.9, 42mM NaCl, 5mM KCl, 2mM MgCl2, 1mM EGTA) (From personal correspondence with James Priess). Dissected gonads were transferred to poly-lysine treated dish with 80 ul of Happy Buffer and fixed by adding equal volume of 5% formaldehyde in PBST (PBS+0.1% Tween 20) for 30 min. After one wash with PBST, gonads were treated with PBST-Triton (PBST+0.1% Triton) for 10 min, washed with PBST again and emerged in 70% ethanol for 30 min to overnight. Before hybridization, samples were washed with fresh wash buffer (2xSSC +10% formamide) for 5 min. hybridization was performed at 37 °C for 18 hours to overnight in hybridization buffer (900 ul Stellaris RNA FISH Hybridization Buffer+ 100ul formamide) with 10 pmol RNA FISH probes. Samples were washed with wash buffer, once quick wash, one wash for 30 min at 37 °C and two quick washes. Mounting medium with DAPI was added to preserve the signal. Confocal images were taken with an Andor Dragonfly Spinning Disk confocal microscope and processed with Fusion and Imaris.

### DNA FISH

Same to RNA FISH, gonads were dissected, fixed and washed with PBST and treated with 70% ethanol. Then, samples were washed with wash buffer three times, one at room temperature for 5 min, one at 95 °C for 3 min, and one at 60 °C for 20 min. Hybridization was performed in hybridization buffer (700 ul Stellaris RNA FISH Hybridization Buffer + 300 ul formamide + primary probes (final 10 pmol) + detection probe (final 10 pmol)) at 95 °C for 5 min and then transferred to 37 °C for 3 hours to overnight. After hybridization, samples were wash with 2xSSC for 20 min at 60 °C, and then 2xSSCT (2xSSC + 0.3% Triton X-100) for 5 min at 60 °C and another 20 min at 60 °C. After another wash with 2xSSCT for 5min at room temperature, samples were preserved in the mounting medium with DAPI. Confocal images were taken with an Andor Dragonfly Spinning Disk confocal microscope and processed with Fusion and Imaris. Primary probes of DNA FISH were picked from the oligo lists generated by OligoMiner (Beliveau et al., 2018).

### RNAi Experiments

Synchronous L1 worms of the reporter strain were plated on NGM plates for 48 hours. Then the worms were collected and washed with M9. About 100 worms were plated on every IPTG plate with the *gfp* RNAi food. After 24 hours, worms were dissected for the FISH experiment. RNA FISH and DNA FISH were performed as described above.

## Data availability

Small RNA sequencing data is available in NCBI (PRJNA874806).

## Author Contributions

Conceptualization, Y.D., and C.C.M.; Investigation, Y.D. and H.O..; Methodology, Y.D., H. O., T.I., and C.C.M.; Data analysis, Y.D.; Writing-Review & Editing, Y.D., and C.C.M.; Supervision, C.C.M.

## Acknowledgement

We thank members of Mello and Ambros labs for discussions; James Priess (Fred Hutchinson Cancer Center) for sharing the receipt of happy buffer and imaging experiences; Weifeng Gu (University of California, Riverside) for providing the PIR-1 protein for small RNA cloning; Ahmet Ozturk for building the small RNA analysis pipeline; Darryl Conte for critical comments and edits on the manuscript; RNA Therapeutics Institute for offering the Nextseq 550 sequencing machine. The work was supported by NIH funding (GM058800 and HD078253) to C.C.M. C.C.M. is a Howard Hughes Medical Institute Investigator.

## Supplementary Figures

**Figure S1.**
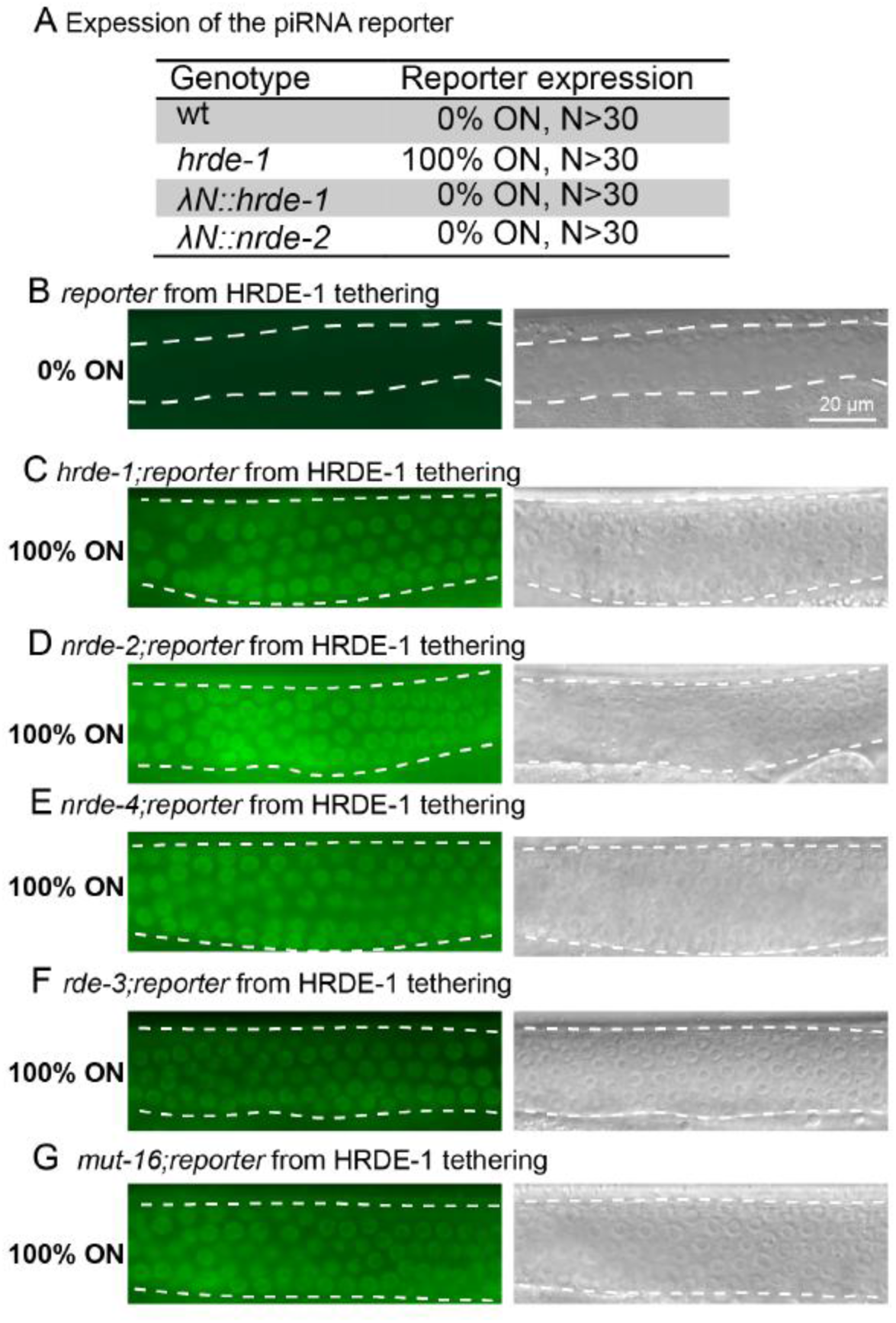
(A) Table shows the score of piRNA reporter expression in wild type and mutants. (B-G) Representative fluorescence (left panels) and DIC (right) images showing the requirement of inherited silencing triggered by HRDE-1 tethering (For Figure 1G). Percentage of worms with reporter ON were indicated, N>30.

**Figure S2.**
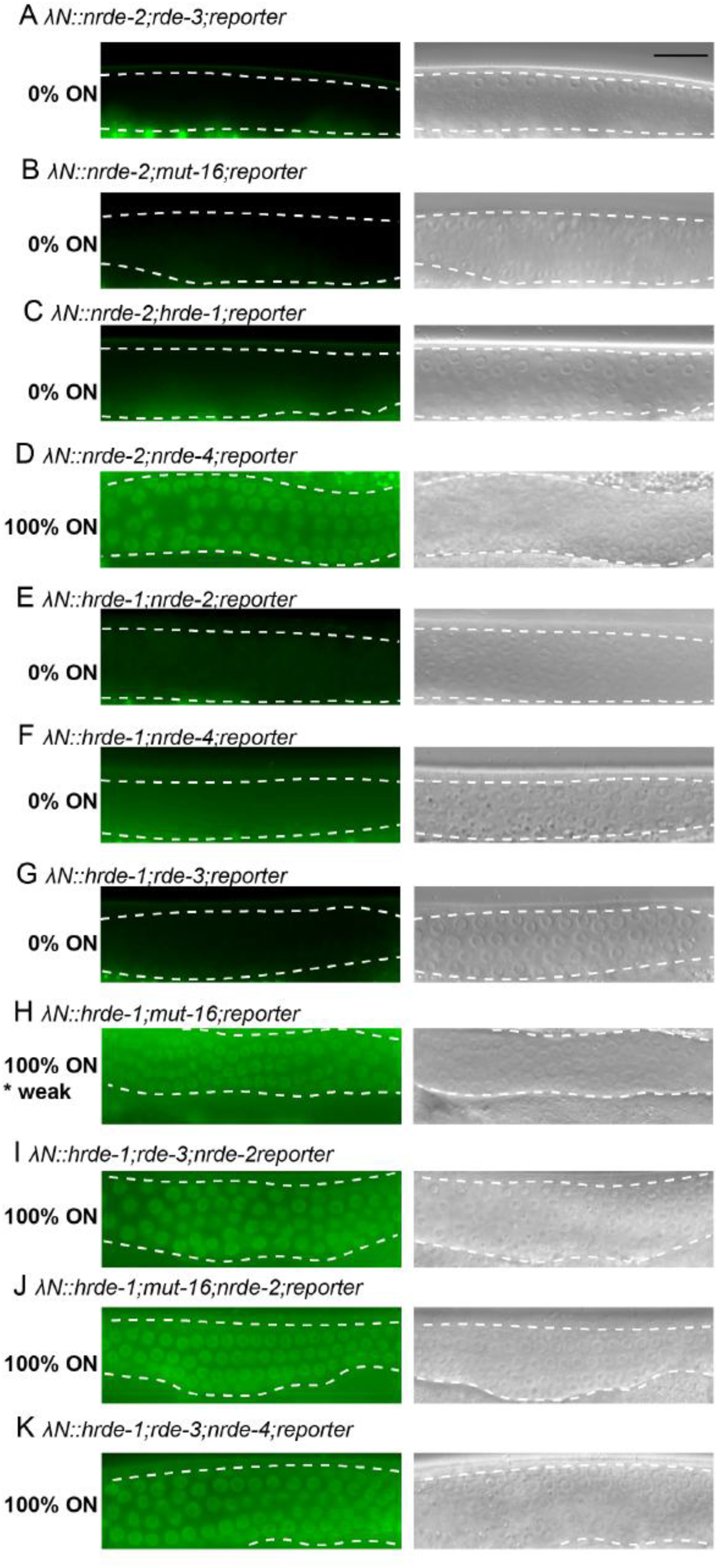
(A-D) Representative fluorescence (left panels) and DIC (right) images showing the expression of boxB reporter in the presence of NRDE-2 tethering in corresponding mutants (For Figure 2E). Percentage of worms with reporter ON were indicated, N>30. (E-K) Representative fluorescence (left panels) and DIC (right) images showing the expression of boxB reporter in the presence of HRDE-1 tethering in corresponding mutants (For Figure 2F). N>30.

**Figure S3.**
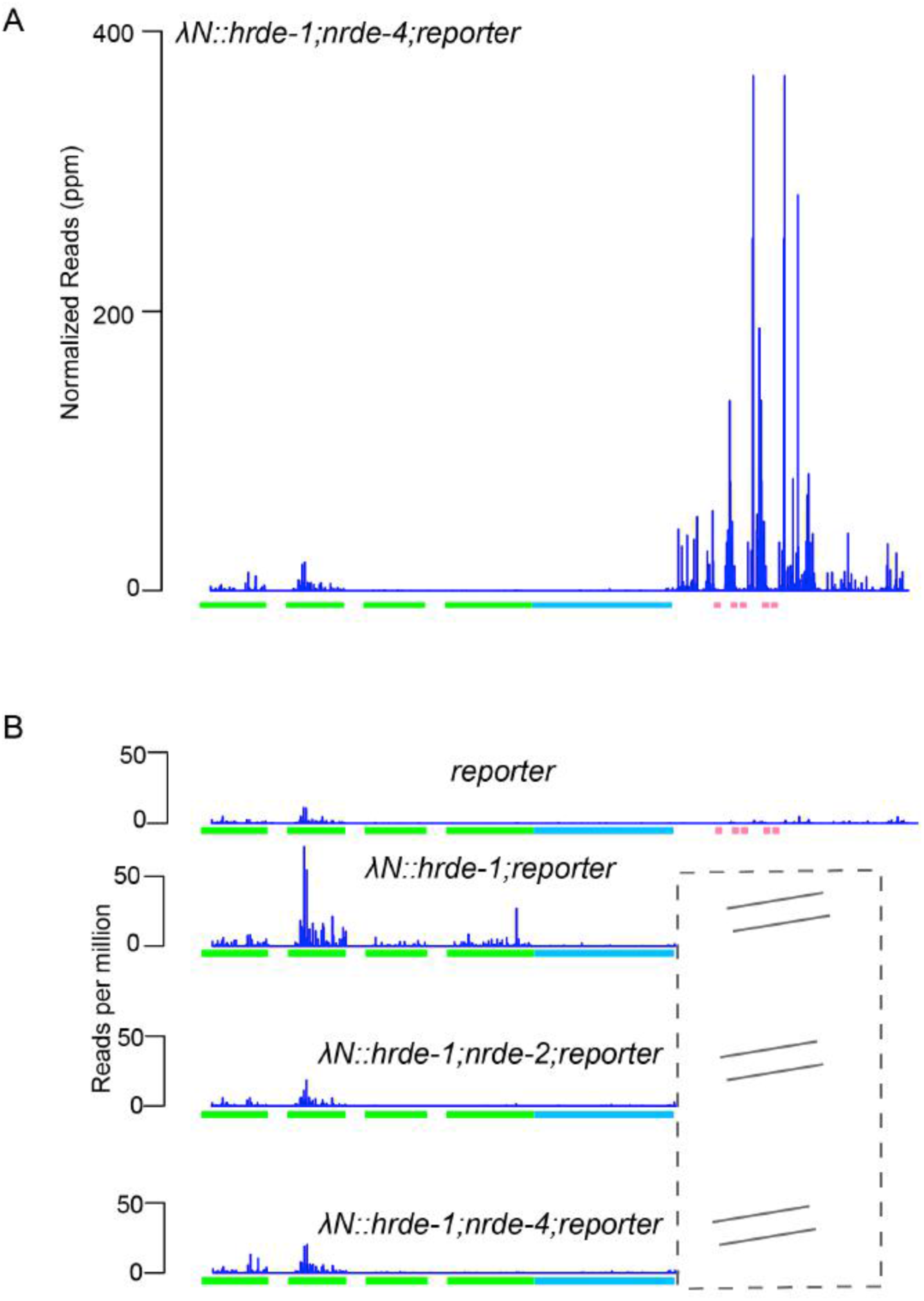
(A) Plot showing antisense small RNA reads (per million total reads) mapping to the reporter (indicated blow the plot) in the presence of HRDE-1 tethering in the *nrde-4* mutant. (B) Plot showing antisense small RNA reads (per million total reads) mapping to the reporter in the presence of HRDE-1 tethering (corresponding to Figure 3F, 3G and S3A). Reads mapping to the BoxB region and 3’ UTR were excluded.

**Figure S4.**
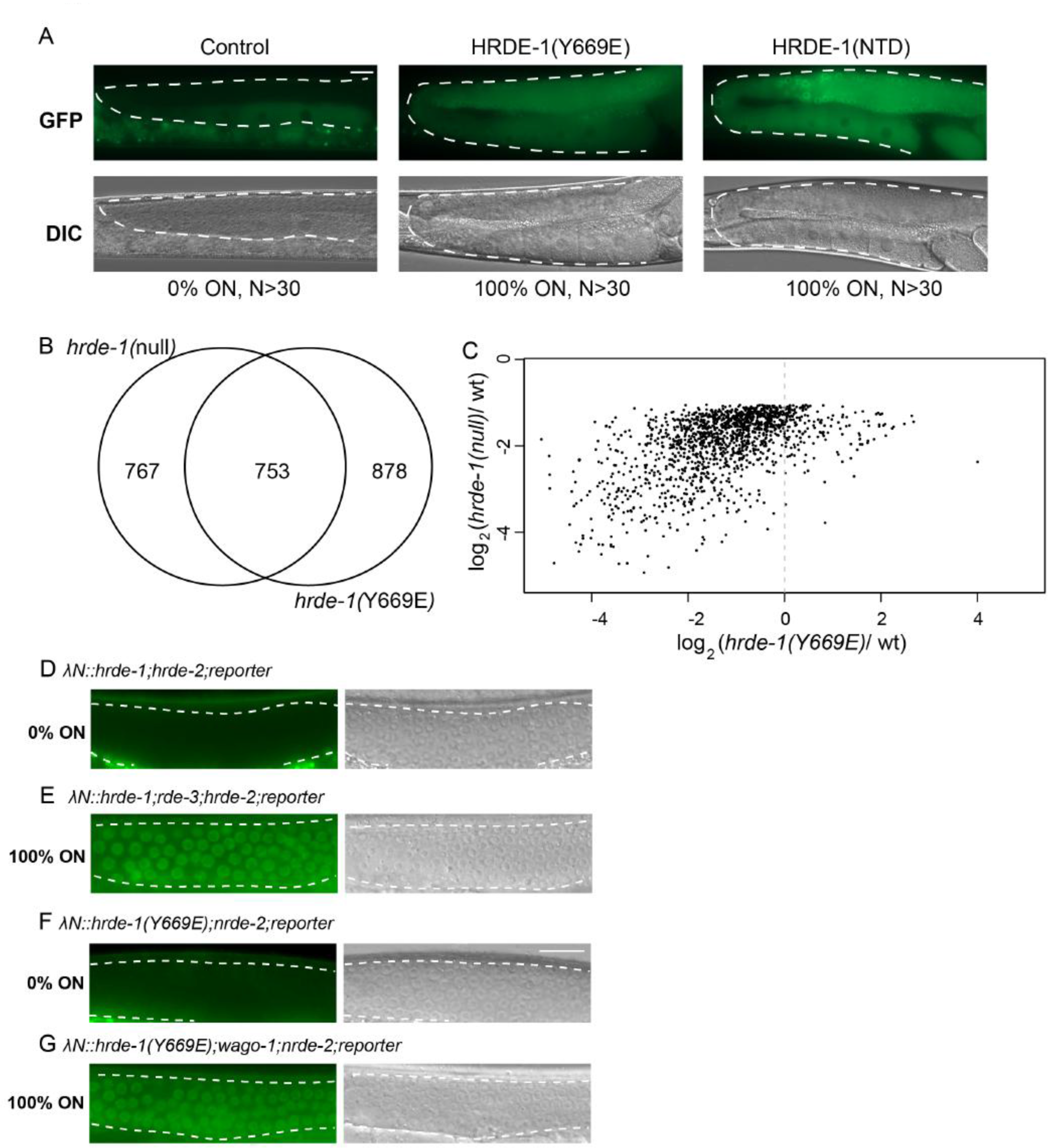
(A) Representative fluorescence (upper panels) and DIC (lower) images showing the expression of the piRNA reporter in wild type, *hrde-1(Y669E)* and *hrde-1(NTD)*. (B) Ven diagram showing the overlap of genes whose antisense small RNA decreased (2-fold and P<0.05) between *hrde-1(null)* and *hrde-1(Y669E)*. (C) Plot showing the ratios of antisense small RNA level in *hrde-1(null)* versus wild type and *hrde-1(Y669E)* versus wild type. Only of genes whose small RNA level decreased (2-fold and P<0.05) in *hrde-1(null)* were ploted. (D-G) Representative fluorescence (left panels) and DIC (right) images showing the expression of boxB reporter in the presence of HRDE-1 or HRDE-1(Y669E) tethering in corresponding mutants. Percentage of worms with reporter ON were indicated, N>30.

**Figure S5.**
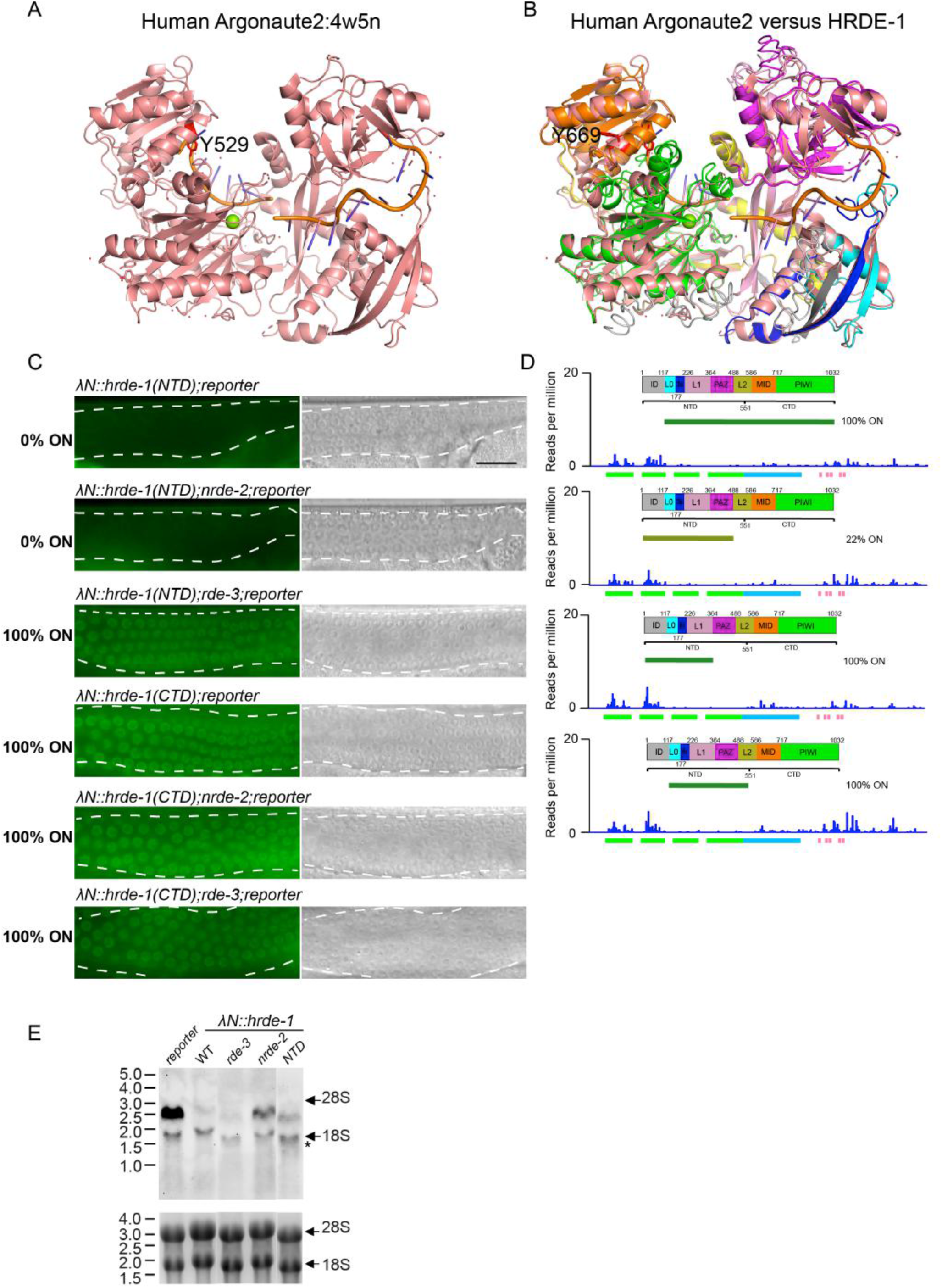
(A) Three-dimensional structure of human Argonaute 2 protein (PDB: 4w5n). hAgo2 is colored in pink and small RNA is colored in yellow. Image was generated by PyMOL. The phosphorylation site Y529 was indicated and colored in red. (B) Overlayed structures of hAgo2 and HRDE-1 (predicted by iTASSER). Two structures were aligned and illustrated by PyMOL. HRDE-1 domains were colored as Figure 5A. the predicted phosphorylation site Y669 of HRDE-1 was indicated and colored in red (C) Representative fluorescence (left panels) and DIC (right) images showing the expression of boxB reporter in the presence of HRDE-1 NTD or CTD tethering in wild and mutants. Scale bar is 20 um. (D) Plot showing antisense small RNA reads (per million total reads) mapping to the reporter (indicated blow the plot) in HRDE-1 truncation tethering. Scheme of the corresponding truncation is indicated in every plot. (E) Northern blot showing the abundance of the reporter RNA in the presence of HRDE-1 tethering in wild type and mutants. 18S and 28S RNA were used as internal control (lower panel). Arrows in the upper panel point to the position of 18S and 26S RNA. * non-specific RNA bands.

**Figure S6.**
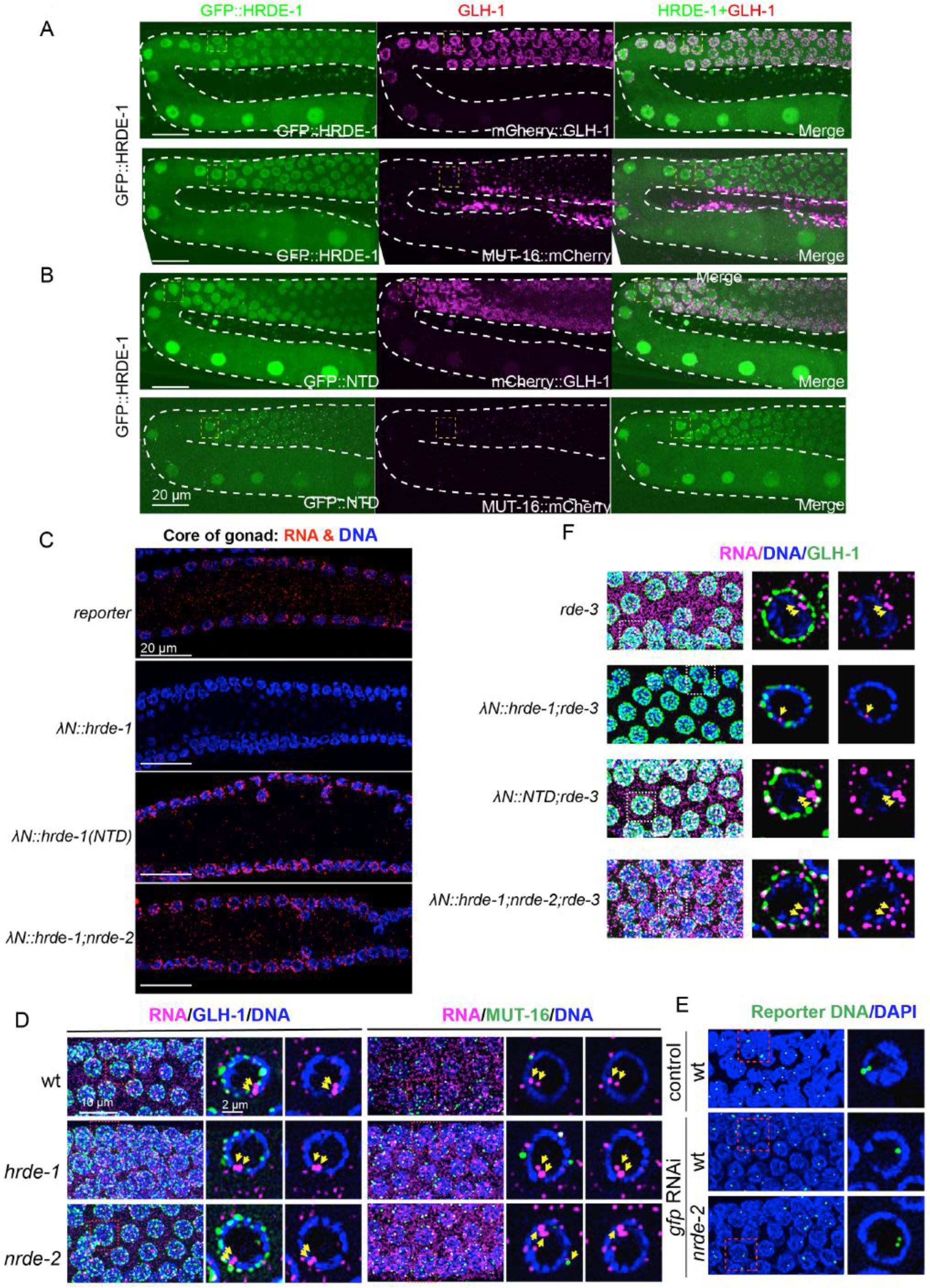
(A) Confocal images showing the localization of GFP::HRDE-1 with mCherry::GLH-1 (upper panel) or MUT-16:mCherry (lower panel) corresponding to Figure 6A and 6B. Gonad is indicated inside the dashed lines. Scale bar is 20 um. (B) Confocal images showing the localization of GFP::HRDE-1(NTD) with mCherry::GLH-1 (upper panel) or MUT-16:mCherry (lower panel) corresponding to Figure 6C and 6D. (C) Confocal images showing the RNA FISH signal in the germline cytoplasm in the absence and presence of HRDE-1 tethering. Only middle slices of z stacks were projected. DNA from DAPI staining was shown in blue. (D) Confocal images of RNA FISH experiments showing the localization of reporter RNA in wild type, *hrde-1* and *nrde-2* mutants with mCherry::GLH-1 (left) or MUT-16::mCherry (right). Yellow arrows point to the nuclear RNA signals. (E) Confocal images of DNA FISH experiments showing the localization of the reporter DNA loci in the absence or presence of *gfp* RNAi. Green, DNA FISH signal; blue, DAPI. A projected view of a segment of a representative germline is shown to the left and the nucleus bounded by a dashed box is shown as a single-focal-plane image to the right. (F) Confocal images of RNA FISH experiment showing the localization of reporter RNA in the absence or presence of HRDE-1 tethering in *rde-3* mutants. Magenta, RNA FISH signal; green, mCherry::GLH-1; blue, DAPI. A projected view of a segment of a representative germline is shown to the left and the nucleus bounded by a dashed box is shown as a single-focal-plane image to the right. Yellow arrows point to the nuclear RNA signals.

**Figure S7.**
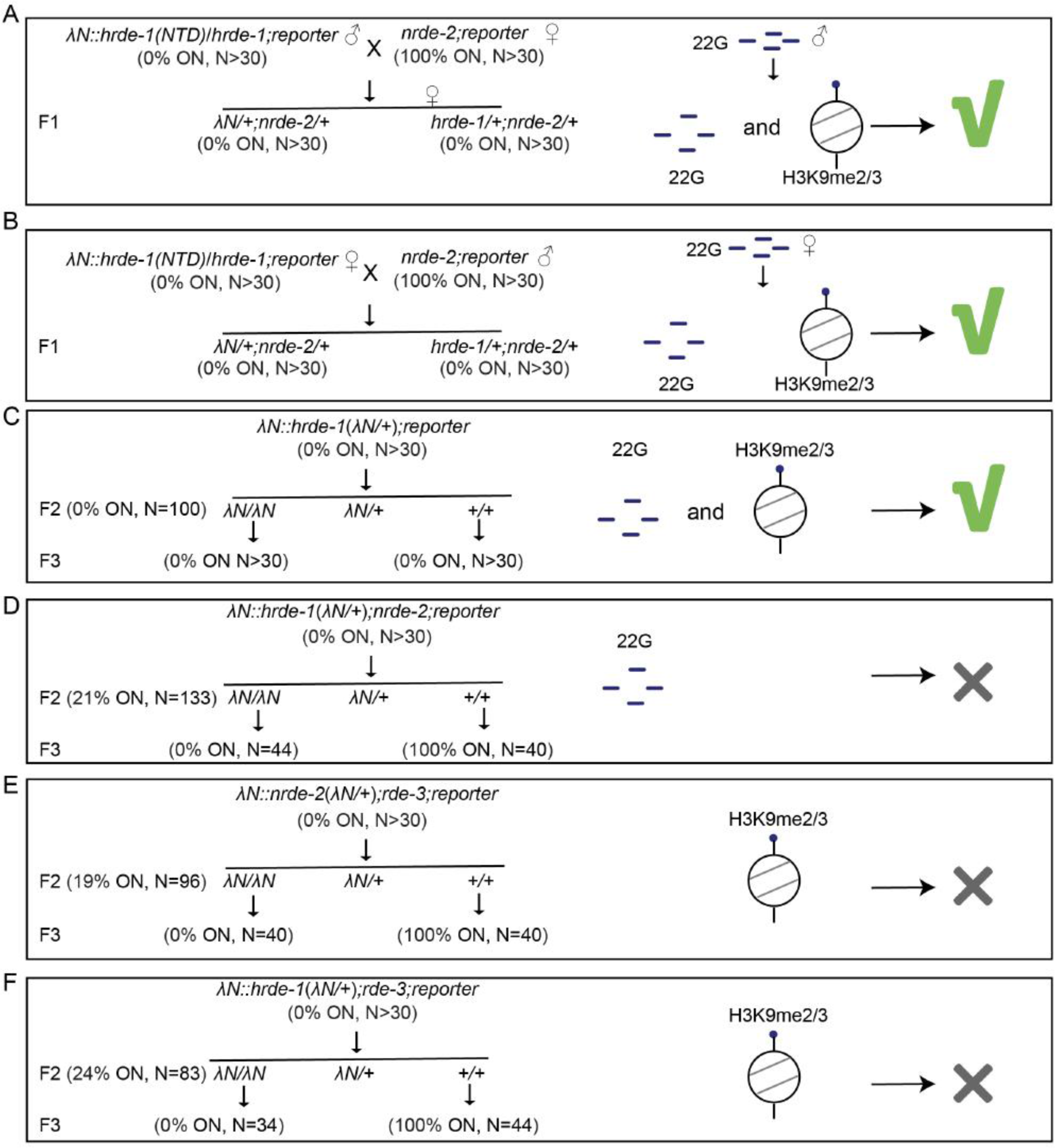
Genetics analysis of the role of 22G and heterochromatin in transgenerational silencing. The mating process was shown to the left panel and the scheme of 22G and heterochromatin involved in the process to the right. Green check mark 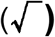 indicates the silencing memory was transmitted to the progeny while **X** is for losing silencing memory.

